# Macrophage inflammatory and regenerative response periodicity is programmed by cell cycle and chromatin state

**DOI:** 10.1101/2021.06.24.449850

**Authors:** Bence Daniel, Julia A. Belk, Stefanie L. Meier, Andy Y. Chen, Katalin Sandor, Yanyan Qi, Hugo Kitano, Joshua R. Wheeler, Deshka S. Foster, Michael Januszyk, Michael T. Longaker, Howard Y. Chang, Ansuman T. Satpathy

## Abstract

Cell cycle (CC) is a fundamental biological process with robust, cyclical gene expression programs to facilitate cell division. In the immune system, a productive immune response requires the expansion of pathogen-responsive cell types, but whether CC also confers unique gene expression programs that inform the subsequent immunological response remains unclear. Here we demonstrate that single macrophages adopt different plasticity states in CC, which is a major source of heterogeneity in response to polarizing cytokines. Specifically, macrophage plasticity to interferon gamma (IFNG) is substantially reduced, while interleukin 4 (IL-4) can induce S-G2/M-biased gene expression. Additionally, IL-4 polarization shifts the CC-phase distribution of the population towards G2/M phase, providing a mechanism for reduced IFNG-induced repolarization. Finally, we show that macrophages express tissue remodeling genes in the S-G2/M-phases of CC, that can be also detected *in vivo* during muscle regeneration. Therefore, macrophage inflammatory and regenerative responses are gated by CC in a cyclical phase-dependent manner.

**Highlights:**

- Single-cell chromatin maps reveal heterogeneous macrophage polarization states
- Cell cycle coincides with heterogeneity and alters macrophage plasticity to polarizing cytokines
- Macrophage polarization is a cell cycle phase-dependent immunological process
- S-G2/M-biased gene expression is linked to tissue remodeling and detected in proliferating macrophages during muscle regeneration

## Introduction

Cellular plasticity describes the phenotypic flexibility and responsiveness of a cell type in a changing microenvironment, a feature that is critical for the adaptation to environmental challenges. How plasticity is established in a population of cells is a key question in biology. Interestingly, certain cell types possess the ability to adopt more nuanced phenotypic traits in response to stressors and can also revert from this state, thus being more plastic. In the immune system, this is a particularly important cellular feature, especially at the first line of defense, among the patrolling, long-lived innate immune cell types of blood and tissues.

Macrophages (MF) are innate immune cells with remarkable plasticity. As resident cell of various organs, MFs adopt distinct phenotypes to maintain tissue integrity and resolve infections. They achieve this by quickly adjusting their epigenetic and gene expression programs to the changing microenvironment, a phenomenon called MF polarization [1–4]. In the lung, alveolar MFs respond to infections (e.g. influenza, *Streptococcus Pneumoniae*) and have an important role in surfactant metabolism [4]. Similarly, Kupffer cells of the liver respond to pathogens but also appear to play a central role in metabolizing toxic or carcinogenic compounds [5]. The pleiotropic action of MF subpopulations across tissues indicate the existence of diverse MF plasticity states, tuning their responses at the subpopulation level when they undergo phenotypic polarization upon environmental challenges. Indeed, single-cell studies have begun to reveal MF heterogeneity in multiple tissues and cancer, but no major mechanism has been offered for the formation of the observed heterogeneous phenotypes, and whether certain subpopulations exist in different plasticity states that would affect their polarization capacity [6–8]. Importantly, a common property of MFs is their proliferative potential in the tissue of residence which can be induced by MF (granulocyte) colony stimulating factors (M-CSF and GM-CSF) and the T-helper 2 (Th2)-type cytokine, interleukin 4 (IL-4) resulting in cell cycle (CC) entry [9–13]. MF proliferation is important to replenish the tissue resident pool during both homeostatic and pathological conditions, and has been linked to the resolving phase of inflammation and tissue regeneration [13–17]. However, whether CC influences MF plasticity or polarization capacity has not been determined.

In order to uncover the phenotypic plasticity of MFs, models of classical (exposure to interferon gamma (IFNG) or lipopolysaccharide (LPS) – referred to as M1 MFs) and alternative (exposure to IL-4 or IL-13 – referred to as M2 MFs) MF polarization have become the gold standard approach to understand the molecular principles of MF responses *in vitro* [18–21]. These cellular models uncovered the remarkably dynamic responses of MFs to polarizing cytokines, which can offer a direct measure of their plasticity [22–25]. Although tissue environments harbor a complex molecular milieu and as a result contain a spectrum of MF polarization states, these *in vitro* models proved useful to mimic the most robust MF responses that can also occur *in vivo*. For example, *bona fide* M1 MFs (*Nos2*^+^, *Il1b*^+^, *Tnf*^+^) are present during bacterial or viral infections, while M2 MFs (*Chil3*^+^, *Retnla*^+^, *Arg1*^+^) have been observed in wound healing, helminth infections, and allergic reactions [18, 19, 21]. Therefore, this model is ideal to investigate MF responses to polarizing cytokines to identify fundamental mechanisms that regulate plasticity and might be also translatable to *in vivo* settings.

MF polarization has almost exclusively been studied at the population level [22–28]. Therefore, our view on the transcriptomic and epigenomic programs of MF polarization is hampered by the lack of sub-population level analyses. This apparent gap raises fundamental questions about how MF plasticity is regulated at the single cell level: (1) What are the major determinants of MF plasticity states? (2) Are there cell intrinsic properties that influence plasticity to polarizing signals?

Motivated by these questions, we generated more than 30,000 single bone marrow-derived macrophage transcriptomes (scRNA-seq) and *cis*-regulomes (scATAC-seq) to build a comprehensive genomic atlas of IFNG-induced (M1) and IL-4-induced (M2) macrophage polarization. Using this atlas, we define heterogeneous MF polarization states. We report that CC coincides with heterogeneity and is a major factor that influence MF plasticity during M1 and M2 polarization by sorting MFs from the different phases of CC. Interestingly, MFs lose their plasticity to IFNG in the S-G2/M phases of CC, while IL-4 can induce a specific gene signature at these CC-phases, correlating with phase-biased enhancer activities. We find that CC negatively affects the formation of a chromatin imprint that defines a subpopulation of “memory” MFs. Additionally, CC also limits MF repolarization with IFNG from a M2 state. Finally, we discover a CC-intrinsic tissue remodeling gene signature linked to the S-G2/M phases of CC that can be also detected in proliferating MFs during muscle regeneration. Therefore, our work establishes the connection between CC and MF immune responses, introducing the concept of cyclical immune plasticity, which we propose to be broadly relevant to the cells of the immune system.

## Results

### Single-cell chromatin accessibility landscape of MF polarization

In order to understand MF heterogeneity at the chromatin level, we performed single-cell assay for transposase accessible chromatin using sequencing (scATAC-seq) of mouse bone marrow-derived resting (unstimulated; M0; CTR), classically-polarized (M1; IFNG), and alternatively-polarized MFs (M2; IL-4) **(Figure 1A)**. In total, we obtained high quality scATAC-seq data from 20,275 single cells from these 3 conditions with a unique fragment count above 1,000 per nuclei and a median read enrichment at transcription start sites (TSSs) of >11 **(Figure S1A and S1B)**. We performed dimensionality reduction using iterative latent semantic indexing (LSI) followed by UMAP visualization, which revealed a clear separation between M0, M1 and M2 MF chromatin states **(Figure 1B)**. Known M2 (*Arg1*) and M1 (*Cxcl9*) polarization marker genes exhibited specific chromatin remodeling in the respective polarization states (quantified by Gene score, see Methods), correlating with bulk gene expression levels **(Figure 1C and D)**. Transcription factor (TF) footprint analysis at polarization-specific TF motifs showed strong footprints at STAT6 and EGR motifs in the M2 condition, while IRF and STAT1 footprints were the strongest in the M1 condition, confirming our polarization model, and recapitulating previously described hallmarks of MF polarization **(Figure S1C)** [23, 29, 30].

**Figure 1.**
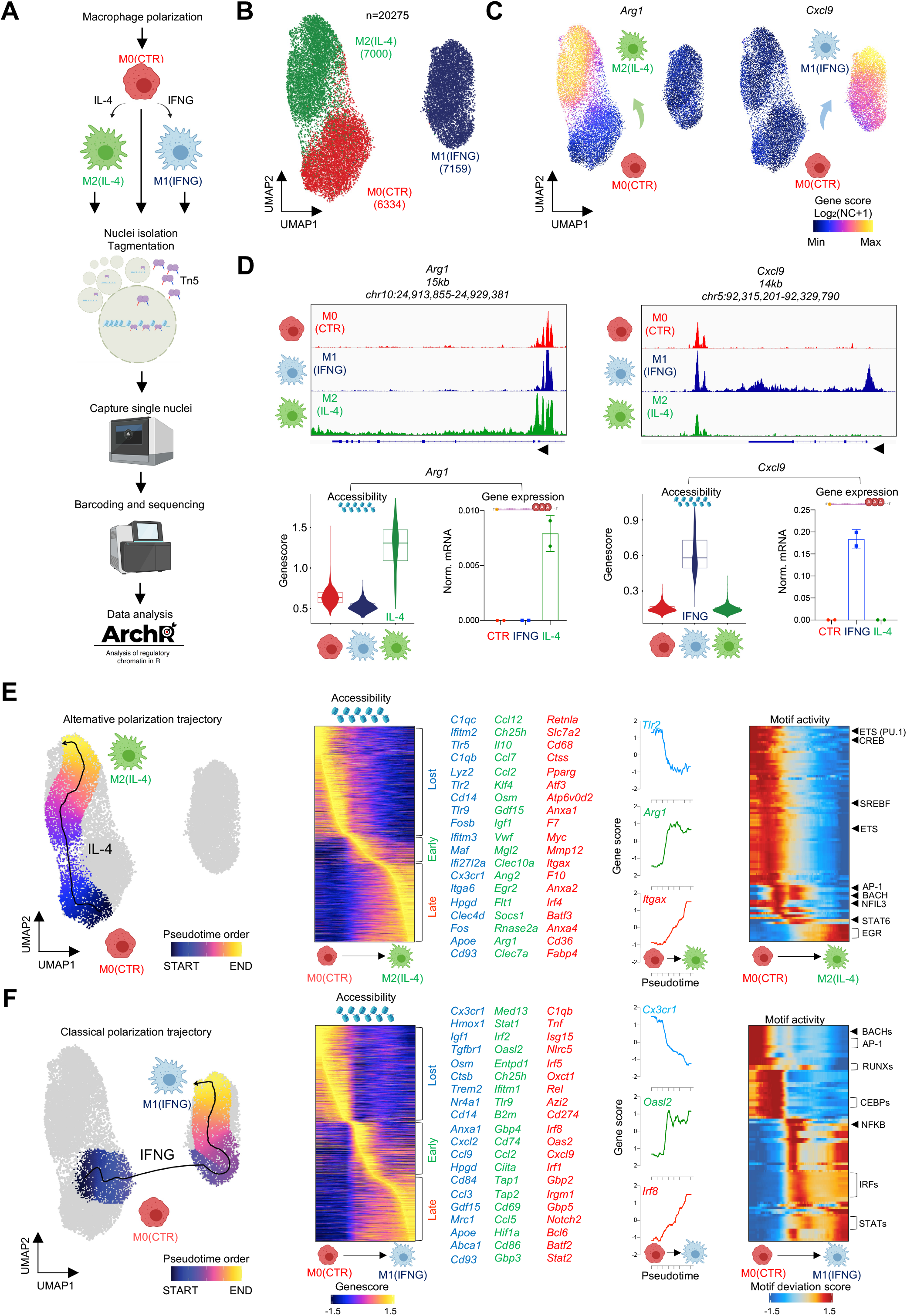
Single cell cis-regulatory program identifies heterogeneous macrophage subsets during polarization. **(A)** Schematic of the experimental system used. **(B)** UMAP projection of scATAC-seq results on polarized macrophages. **(C)** UMAP projection of gene score (accessibility) values for *Arg1* and *Cxcl9*. **(D)** Genome browser views of scATAC-seq results in bulk on the *Arg1* and *Cxcl9* loci (top). Bulk gene score values of CTR (red), IFNG (blue) and IL-4 (green) polarized macrophages for *Arg1* and *Cxcl9* along with their mRNA levels as determined by RT-qPCR (bottom). **(E)** UMAP projection of the alternative polarization trajectory on the M0(CTR) – M2(IL-4) transitional states (left). Heatmap of gene scores changing over the polarization trajectory. Genes that lose- (Lost - blue), gain early- (Early - green), or late (Late - red) accessibility are marked and a select set is displayed. *Tlr2*, *Arg1* and *Itgax* gene scores are shown over the pseudotime. ChromVAR transcription factor motif deviation scores over pseudotime on the alternative polarization trajectory. **(F)** Same as panel E; for the Classical polarization trajectory that describes the transitional states between M0(CTR) – M1(IFNG).

We observed a continuum of MF polarization states in the M1 and M2 conditions which prompted us to assess polarization trajectories that can inform phenotypic state transitions [31]. We ordered the single MF chromatin states along a vector that describes the paths of the two main polarization trajectories on the UMAP. First, we reconstructed the M2 polarization trajectory by the nearest-neighbor approach starting from M0 to M2 MFs by sequentially selecting MFs with similar chromatin states (Euclidean distances of single cell chromatin states) [31, 32]. We observed the early and late chromatin remodeling activities of M2 polarization, such as early chromatin closure at repressed genes (e.g., *Tlr2*, *Ifitm2*, *Cd14* and *Hpgd*) and opening around “early” induced genes (e.g., *Arg1*, *Mgl2*, *Egr2* and *Klf4*) [23]. At the later points of the trajectory, we detected chromatin opening around the genes of the late M2 program, including *Retnla*, *Anxa2*, *Mmp12*, and *Pparg* **(Figure 1E; Table S1)** [30]. Importantly, the dynamics of chromatin remodeling at specific genes over the trajectory followed their expression level from a published time course bulk RNA-seq experiment [23]. This result argues that the transitional chromatin states of single MFs, resulting from a 24-hour long M2 polarization can reliably recapitulate the entire *cis*-regulatory/gene expression cascade of MF polarization **(Figure S1D and S1E)**. Motif accessibility analyses over the pseudotime trajectory linked the STAT6 motif to the early chromatin remodeling activities of M2 polarization, while the EGR2 motif was linked to late chromatin remodeling, confirming previous findings **(Figure 1E and S1F; Table S2)** [22, 23, 30].

Next, we performed the trajectory analysis of classical polarization. MF responses to IFNG was more uniform; and MF chromatin states separated more clearly on the UMAP with no transitional states between M0 and M1 MFs **(Figure 1F)**. The trajectory featured the gene set of early chromatin closure (e.g., *Cx3cr1*, *Igf1* and *Cd14*); and early and late chromatin opening, including *bona fide* IFNG-induced genes (Early - e.g., *Irf2*, *Oasl2* and *Stat1*; Late - e.g., *Irf8*, *Irf5* and *Cxcl9*) **(Table S3)** [29]. Motif accessibility analysis suggested the immediate early action of IRF and STAT1 motifs and decreased chromatin accessibility at BACH, AP-1, RUNX and CEBP motifs over the trajectory from M0 to M1 states **(Figure 1F and S1F; Table S4)**. Altogether, this single cell atlas reveals the transitional chromatin state program of M1 and M2 MF polarization and motivated us to further investigate the chromatin structure of heterogeneous MF subsets.

### MF heterogeneity coincides with cell cycle

To identify the main subpopulations of MFs, we clustered the cells and identified two distinct chromatin state clusters in each condition (M0 – C5 and C6; M2 – C1 and C2; M1 – C3 and C4) **(Figure 2A)**. In general, subpopulations of the polarized states did not co-cluster with the clusters of the M0 state. More specifically, no M1 polarized MFs were present in the M0 clusters, while approximately 10% of M2 cells remained in the M0 clusters (C5 and C6) **(Figure 2A)**. Next, we identified the marker gene scores of each cluster (FDR≤0.01, Log_2_ fold change (FC)≥1.25) **(Table S5)**. We observed C1- (n=261) and C2-biased (n=113) gene scores, including several M2 marker genes in the two clusters (C1 - e.g., *Retnla*, *F10*, *F7* and *Abcg1*; C2 - e.g., *Mgl2*, *Igf1*, *Ccl7* and *Ccl2*). Conversely, M1 MFs exhibited C3- (n=483) and C4-biased gene scores (n=317), including *bona fide* M1 marker genes in both clusters (C3 - e.g., *Gbp2*, *Gbp10*, *Ifit3*, *Cd274* and *Cxcl9*; C4 - e.g., *Mmd2*, *Nlrp9b* and *Oas1c*) **(Figure 2B)**. We noticed that cell cycle (CC) gene scores were largely specific to polarized M2 MFs in C2 (e.g., *Hist1h3g*) or M1 MFs in C4 (e.g., *Top2a* and *Ccnf*). Importantly, the gene score values of *Mki67* aligned with the CC gene scores in these clusters, indicating that MFs in C2 and C4 are engaged in CC **(Figure 2C)**.

**Figure 2.**
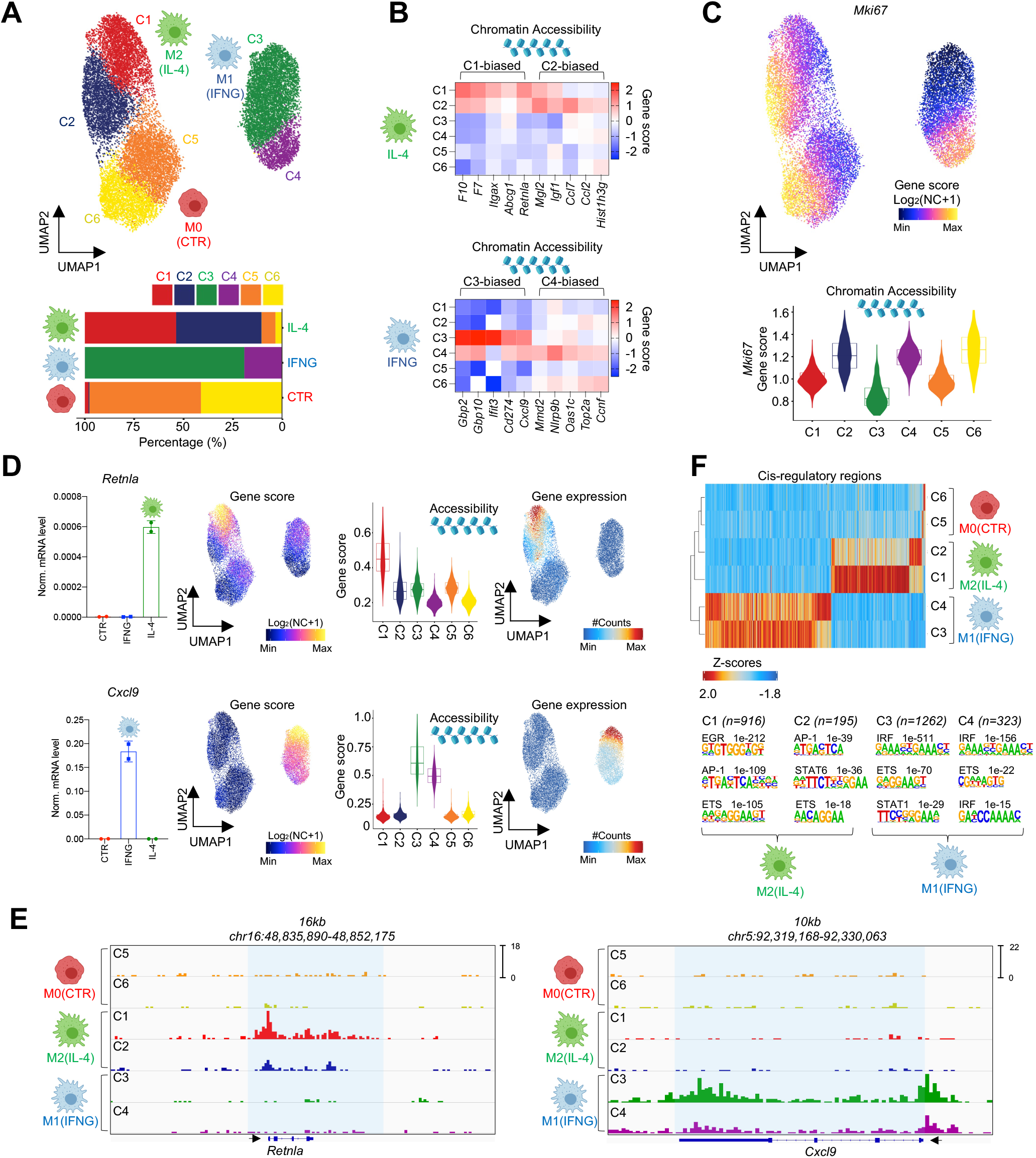
Macrophage heterogeneity coincides with cell cycle markers. **(A)** scATAC UMAP of macrophage polarization colored by the 6 macrophage clusters. Percentage-wise distribution of the clusters across the M0(CTR), M1(IFNG) and M2(IL-4) samples (bottom). **(B)** Heatmap representation of a select set of marker gene scores in the clusters of either M2(IL-4) or M1(IFNG) macrophages. **(C)** UMAP of *Mki67* gene scores. Violin plot of *Mki67* gene scores in the clusters. **(D)** Bar graphs depict bulk mRNA levels of *Retnla* and *Cxcl9* (left). UMAPs and violin plots show the gene score values (log_2_ normalized counts+1) (scATAC-seq) for the two genes (middle). UMAPs of gene integration scores (gene expression - scRNA-seq), # - normalized. **(E)** Genome browser views of scATAC-seq signal in the 6 clusters on the *Retnla* and *Cxcl9* loci. **(F)** Peak score heatmap of differentially accessible cis-regulatory regions in the clusters (top). Homer *de novo* motif search results on the cluster-specific peaks. The number of regions in each cluster and the p-values for the enriched motifs are shown (bottom).

To link transcriptional activity to the observed chromatin changes, we performed single cell RNA-seq in M0, M1 and M2 MFs, and identified the differentially expressed genes between the M0 – M2 (Induced: 214, Repressed 147) and M0 – M1 (Induced: 494, Repressed: 212) conditions (FDR≤0.01, Log_2_ FC≥0.25) **(Figure S2A and S2B; Table S6)**. We performed constrained integration of single cell chromatin and transcriptomic profiles of the different polarization states, thereby limiting the search space and enhancing the efficiency of the integrative method [32]. As a result, we generated a gene integration matrix that contains pseudo scRNA-seq expression values linked to each cell in the scATAC-seq space, which recapitulated our observations on cluster-biased chromatin accessibility at the mRNA level **(Figure S2C)**. For example, the expression of CC genes appeared to be specific to C2 of M2 and C4 of M1 polarized MFs (e.g., *Cenpt*, *Top2a, Kif4* and *Cdk1*). These clusters of cells in CC (hereafter referred to as cell cycle (CC)-clusters) exhibited reduced expression profiles for specific M1 (e.g., *Cxcl9* and *Gbp2*) and M2 genes (e.g., *Retnla* and *Egr2*) **(Figure 2D, E and S2C)**; however, we also observed genes with CC-cluster-biased expression, such as *Mgl2* in M2 polarized MFs (C2) **(Figure S2C)**. Collectively, these results show that MF heterogeneity coincides with CC and suggests that MF plasticity to polarization cytokines is influenced by CC.

### The *cis*-regulatory landscape of M1 and M2 MF polarization is constrained by cell cycle

After we defined the chromatin states of polarization-specific genes and their expression profiles in the subpopulations of M1 and M2 MFs, we turned our attention to identify the distant regulatory regions of the non-coding genome (referred to as *cis*-regulatory elements - CREs). In M2 MFs we identified 916 C1-biased and 195 C2-biased CREs. Analysis of M1 MFs reported 1,262 C3- and 323 C4-biased CREs (FDR ≤0.01, Log_2_ FC≥1) **(Figure 2F)**. We observed that in both M1 and M2 MFs, cells in CC-clusters (C2 and C4) showed less pronounced chromatin remodeling events upon polarization compared to non-cycling cells (C1 and C3). However, we also noted a smaller set of polarization-induced CREs that were biased to the CC-clusters. Motif enrichment analyses at the M2-specific CREs identified the EGR2 motif in C1, while the STAT6 motif showed specific enrichment in C2. In M1 MFs, we detected the STAT1 motif exclusively in C3, while the IRF motif was present in both clusters but showed a more significant enrichment in C3 compared to C4 (p-values: C3 - 1e^−511^ versus C4 - 1e^−156^). Single cell chromatin accessibility analyses of these motifs further supported these findings **(Figure S2D)**.

According to these results, the subset of non-cycling MFs showed the highest level of chromatin remodeling potential in both polarization models. Interestingly, while both the STAT1 and IRF motifs are largely specific to non-cycling M1 MF CREs, the binding motif of STAT6 appears to show specific enrichment in the CC-cluster of M2 MFs, whereas the EGR motif is specifically enriched in the non-cycling MF cluster. These results suggest that STAT6 and EGR2 act in different MF subpopulations after 24 hours of IL-4 polarization, in agreement with a recent study that reported spatial and temporal separation of the binding sites of the two TFs in M2 MFs [30]. Additionally, these results imply that the main TFs of M1 (STAT1 and IRFs) and M2 (EGR2) MFs might lose some of their functions in CC, but STAT6 might be able to retain its transcriptional activity.

### Cell cycle limits the expression of the two key transcription factors of MF polarization

We next asked whether CC alters the plasticity of MFs to polarization signals. First, we took a predictive approach and used our gene integration matrix to assign specific CC stages (G1, S and G2/M) to each cell in the scATAC-seq space using a CC scoring algorithm **(Figure 3A)** [33]. This analysis showed that ~80% of the cells in C2 (M2) and more than 95% of cells in C6 (M0) and C4 (M1) were in CC **(Figure S3A)**. Utilizing the gene integration matrix, we performed differential gene expression analyses (FDR≤0.01, FC≥1.3) between CC stages using the marker genes of M2(IL-4) and M1(IFNG) states that we previously defined **(Figure S2B, Table S6)**. Comparison of G1 to G2/M yielded the largest gene lists in both polarization settings, suggesting that gene expression is largely biased towards the G1-phase (M1: 38 G1-biased genes; M2: 33 G1-biased genes) **(Figure 3B)**. Interestingly, the M2 gene expression program showed more S- (n=6; e.g., *Atpv0d2* and *Anxa1*) and G2/M-biased (n=11; e.g., *Mgl2* and *Gatm*) genes over the G1-phase as compared to the M1 program, suggesting that IL-4 might be able to initiate more specific polarization programs in CC **(Figure S3B)**. The M1 program appeared to be more sensitive to CC and we found a negligible number of genes with biased expression in the later stages of CC (S vs. G1 – n=2; S vs. G2/M – n=1; G2/M vs. S – n=2; G2/M vs. G1 – n=2). Importantly, the G1-biased gene program resulted in a largely overlapping gene set from the G1-S and G1-G2/M comparisons, supporting the idea that IFNG response is severely reduced in the S-G2/M phases of CC **(Figure 3B and S3B)**. Next, we examined the gene sets of the G1-G2/M comparisons in the two models and found attenuated expression of the typical polarization marker genes in the G2/M phase (M2(IL-4): *Anxa1*, *Evl*, *Egr2*, *Atf3* and *Atp6v0a1*; M1(IFNG): *Slfn5*, *Ifit2*, *Ifit1*, *Irf8* and *Ifitm2*) **(Figure 3B)**. We noticed that the TFs, *Egr2* and *Irf8* exhibited G1-biased expression and chromatin accessibility profiles in M2 or M1 polarized MFs, respectively **(Figure S3C and S3D)**. EGR2 is a critical regulator of M2 MF, while IRF8 orchestrates significant parts of the M1 MF polarization program, indicating a dysfunctional polarization program in the later phases of CC [30, 34].

**Figure 3.**
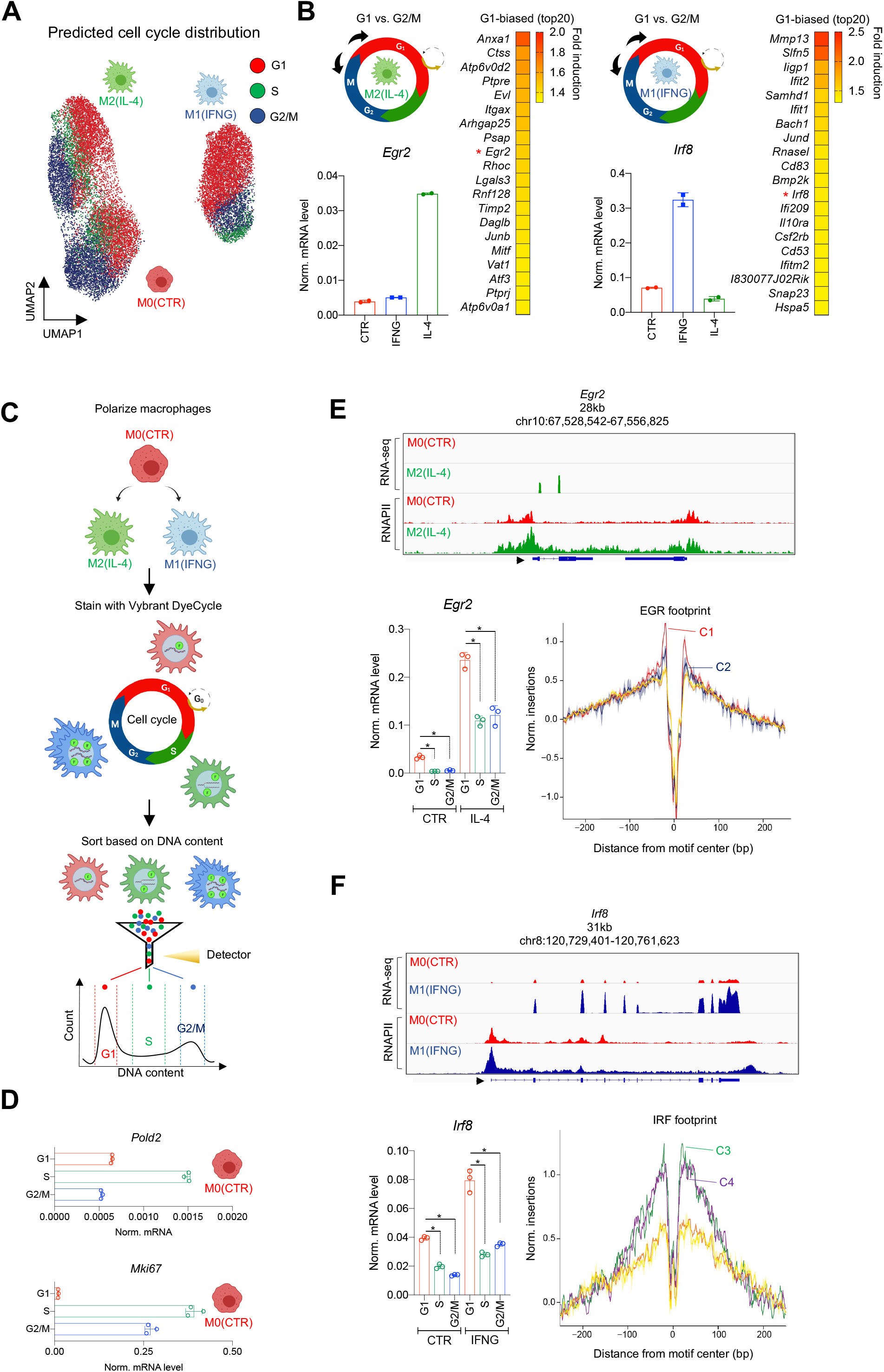
Cell cycle limits the expression of Egr2 and Irf8 during polarization. **(A)** UMAP of cell cycle scores in the polarized macrophage populations. **(B)** Differential gene expression analysis of G1 and G2/M predicted cells from the polarized states (scheme). Heatmap of genes exhibiting G1-biased expression in M2(IL-4) (left) or M1(IFNG) (right). *Egr2* and *Irf8* transcription factors are marked by red asterisks and their bulk expression level is validated by RT-qPCR. **(C)** Scheme of cell cycle sorting. **(D)** mRNA levels of *Pold3* and *Mki67* measured by RT-qPCR in cell cycle phase-sorted macrophages. **(E)** Genome browser view of bulk RNA-seq and RNAPII ChIP-seq results on the *Egr2* locus in M0(CTR) and M2(IL-4) macrophages. mRNA level of *Egr2* in cell cycle phases, significant changes were determined by two tailed, unpaired t-test at p<0.05 (n=3). Shown are means with SDs. EGR transcription factor footprints in the 6 scATAC clusters. **(F)** Same as panel E; for *Irf8*.

To experimentally test these observations, we used fluorescence-activated cell sorting (FACS) to quantify the CC distribution of M0 MFs with a DNA labeling dye (Vybrant DyeCycle) **(Figure 3C)**. This experiment reinforced the notion that MFs can be detected in different CC stages, in agreement with a previous study [35]. Specifically, we detected ~73% of the population in G0/G1 (referred to as G1), ~12% in S and ~6.6% in the G2/M phase of CC **(Figure S3E)**. Next, we sorted F4/80^+^ M0 MFs from the different phases of CC and performed gene expression measurements by real time quantitative PCR (RT-qPCR). Reassuringly, the expression of the S-phase-specific *Pold2* (DNA-polymerase delta complex member required for genome replication) and the S-G2/M-specific *Mki67* genes validated the purity of our sorted populations **(Figure 3D)** [36]. Then, we sorted M1 and M2 MFs from CC-phases and measured the expression of *Egr2* and *Irf8* by RT-qPCR, which are readily induced by either IL-4 or IFNG, respectively (assessed by bulk RNA-seq and RNAPIIpS2 ChIP-seq datasets) **(Figure 3E and F)** [29]. As expected, based on our predictions, both *Egr2* and *Irf8* were sensitive to CC; and displayed G1-biased expression in the M0 condition. Additionally, M2 polarization rapidly induced the level of *Egr2* in a G1-biased manner, while M1 polarization resulted in a similar, G1-biased expression profile for *Irf8* **(Figure 3E and F)**. Footprint analyses of the two TF motifs reported C1-biased EGR, while C3-biased IRF footprints corresponding to non-cycling MFs, supporting the gene expression results **(Figure 3E and F)**. These results might explain the dominance of G1-biased polarization programs, but also raises questions about the existence of S-G2/M phase-specific polarization programs, especially in M2 MFs.

### Cell cycle phase-dependent MF plasticity influences polarization potential

Next, we studied the effects of CC on MF plasticity. We performed bulk RNA-seq experiments on M0, M1, and M2 MFs sorted from CC-phases. To streamline the analysis, we used the top 50 polarization-induced and -repressed genes defined by our scRNA-seq results **(Table S6)**, which contained the core gene expression signatures of both M1 and M2 MF polarization **(Figure 4A and S4A)** [19, 21]. First, using the bulk RNA-seq results, we defined CC-sensitive genes with differential expression profiles between any two CC-phases in each condition, yielding a total of 8700 genes (Benjamini–Hochberg adjusted p-value ≤0.001; FC≥1.3). Then we overlapped this list with our top 50 induced and repressed marker genes of the two polarization models. We found that 74% of the M2 gene expression program was sensitive to CC (74/100 genes). Namely, 66% of the induced genes and 82% of the repressed genes exhibited CC phase-dependent expression. Similarly, 76% of the core M1 polarization program appeared to be CC-sensitive (76/100 genes); 84% of the induced genes and 68% of the repressed genes displayed CC phase-biased expression **(Figure S4B)**.

**Figure 4.**
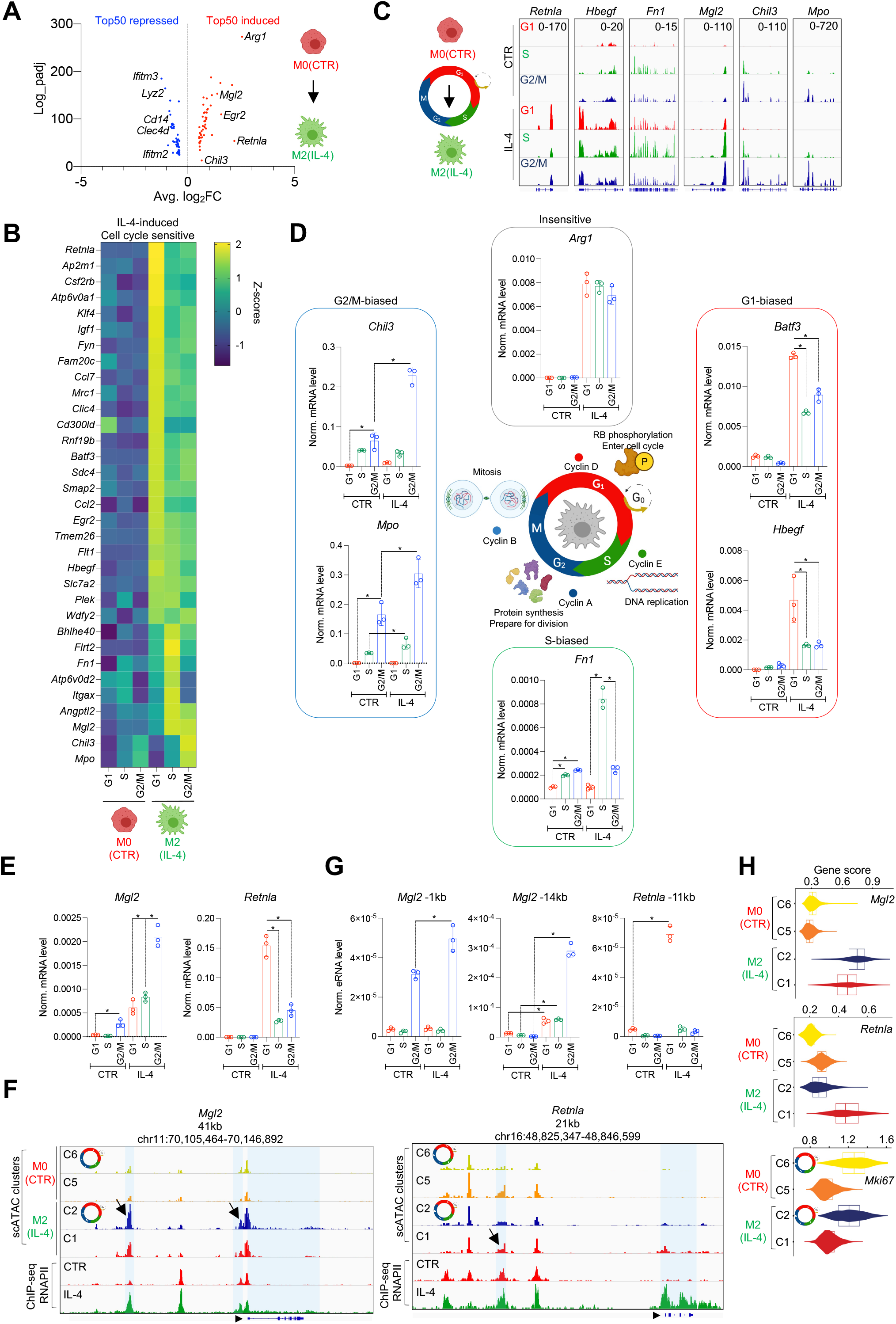
Cell cycle phase specifies macrophage plasticity to polarization signals. **(A)** Volcano plot of the top 50 differentially expressed gene upon M2(IL-4) polarization determined by scRNA-seq. **(B)** Heatmap of cell cycle phase sensitive, IL-4-induced genes determined by bulk RNA-seq. **(C)** Genome browser snapshots on a select set of genes exhibiting phase-biased expression. **(D)** Validation of cell cycle phase-biased gene expression by RT-qPCR. **(E)** Validation of the phase-biased expression of *Mgl2* and *Retnla* by RT-qPCR. **(F)** Genome browser views on the *Mgl2* and *Retnla* loci showing scATAC-seq signal in the clusters of M0(CTR) and M2(IL-4) macrophages along with bulk RNAPII ChIP-seq results in the same conditions. **(G)** Enhancer RNA measurements by RT-qPCR on the *Mgl2* and *Retnla* loci. **(H)** Violin plots depict the gene score values of *Mki67*, *Mgl2* and *Retnla* in the 4 clusters of M0(CTR) and M2(IL-4) macrophages. On the bar graphs, significant changes were determined by two tailed, unpaired t-test at p<0.05 (n=3). Shown are means with SDs.

Next, we analyzed CC phase-biased expression among IL-4-induced genes. We found genes with G1- (48%, 24/50) and S-G2/M-biased expression (18%, 9/50). In both groups, we detected *bona fide* M2 MF marker genes such as the G1-biased *Retnla*, *Atp6v0a1*, *Batf3*, *Hbegf* and *Egr2*, and the S-G2/M-biased *Bhlhe40*, *Fn1*, *Mgl2*, *Chil3* and *Mpo* **(Figure 4B and C, Table S9)**. In contrast to the M2 program, the vast majority of IFNG-induced gene expression circuit (82% - 41/50 genes) exhibited G1-biased expression (e.g., *Cxcl9*, *Ifi44*, *Gbp4* and *Irf8*) and only 3 genes showed S-G2/M-biased expression (*Ccl12*, *Apobec3* and *Pnp*) **(Figure S4B and C, Table S9)**. Among the 50 induced genes in the two polarization models, we found 17 IL-4- (e.g., *Arg1*, *Ptpre*, *Prkcd*, *Gatm* and *Cblb*) and 6 IFNG-induced (e.g., *Cxcl10*, *Gbp5*, *Irf1* and *Fam26f*) but CC-insensitive genes **(Figure S4F and G)**.

Repressed genes exhibited strong phase-biased expression in the M0 condition in both polarization models. Among the IL-4 repressed genes, 62% showed G1-biased expression (e.g., *Cd14*, *Ifitm2* and *Clec4d*) and 25% displayed S-G2/M-biased expression (e.g., *Cx3cr1*, *Spp1* and *Ifi27l2a*) in the M0 condition. In the group of IFNG-repressed genes, 44% had G1-biased expression (e.g., *Ifngr1*, *Plin2* and *C5ar1*) and 24% exhibited S-G2/M-biased expression, including genes that are required for replication, in agreement with the finding that IFNG triggers CC arrest at the G1-S border in MFs (e.g., *Rps28*, *Slbp* and *Gmnn*) **(Figure S4E)** [37]. Therefore, repression occurs by silencing phase-biased gene expression in the M0 state.

Due to our finding that IL-4 can specifically induce gene expression in the S-G2M-phases of CC, we focused on the M2 program and validated our RNA-seq results by RT-qPCR (Insensitive - *Arg1*; G1-biased - *Batf3* and *Hbegf;* S-biased - *Fn1*; G2/M-biased - *Mpo* and *Chil3*) **(Figure 4D)**. Importantly, all of these genes reproduced the CC-phase-biased expression patterns that we detected with RNA-seq. Collectively, our findings show that the majority of the core MF polarization program is CC phase sensitive. Surprisingly, IFNG-induced gene expression is strictly restricted to the G1-phase, whereas IL-4 can launch specific parts of the M2 polarization program in the S-G2/M phases of CC.

### Cell cycle phase-biased expression of *Mgl2* and *Retnla* associates with phase-biased enhancer activities

Next, we set out to study the potential mechanism of phase-biased gene expression. We hypothesized that phase-biased enhancers might drive gene expression based on our observations of biased chromatin remodeling activities in cycling and non-cycling MF clusters **(Figure 2F)**. We focused on two major components of the M2-induced polarization program, *Mgl2* and *Retnla* as demonstrating either G2/M- or G1-biased IL-4-induced expression patterns (RNA-seq – Figure 4B), respectively. Resistin-like alpha (*Retnla*) is one of the most widely accepted markers of M2 polarization in both *in vitro* and *in vivo* conditions. It is a secreted protein exhibiting robust induction in IL-4/-13-induced M2 MFs during multicellular parasite infections (nematodes, helminths) and allergic reactions. It has chemotactic activities towards eosinophils that is speculated to have roles in anti-parasitic defense mechanisms [38]. Macrophage galactose-type C-type lectin (*Mgl2*) is a pattern recognition receptor recognizing glycan structures and implicated in antigen uptake and presentation. Importantly, *Mgl2* is also a widely known M2 marker gene and is induced by Th2-type cytokines during helminth infections and asthma [39, 40]. First, we validated the CC-phase-biased expression of both genes, which confirmed the RNA-seq results **(Figure 4E)**. Second, we inspected their genomic neighborhood, searching for open chromatin regions (scATAC-seq) that align with IL-4-induced RNAPII ChIP-seq signals to identify putative enhancers **(Figure 4F)**. Next, we identified CC-clusters (C6 and C2), according to the clusters defined in Figure 2A and based on *Mki67* accessibility. In agreement with this and the gene expression results, *Mgl2* displayed C2-biased accessibility and *Retnla* showed C1-biased accessibility **(Figure 4F and H)**.

In the *Mgl2* locus, we identified potential enhancers with C2-biased, IL-4-induced accessibility and RNAPII recruitment. Enhancer RNA (eRNA) expression is one of the best markers of enhancer activity; thus, we measured eRNAs at two putative enhancers located −1kb and −14kb from *Mgl2* by RT-qPCR [41, 42]. We detected G2/M-phase-dependent enhancer activity in the M0 state at the −1kb enhancer, while the −14kb was silent. IL-4 readily induced eRNA production at both enhancers, but the two elements showed striking differences. The −1kb enhancer displayed weak IL-4-induced activity, exclusively in the G2/M-phase (CTR vs IL-4 in G2/M, fold change - FC=1.56), whereas the −14kb region showed strong induction upon IL-4 treatment in all phases (G1-FC=4.1, S-FC=9.2) with superior G2/M-biased activity (FC=96) **(Figure 4G)**.

We also identified a candidate enhancer region (−11kb) at the *Retnla* locus and measured enhancer activity. Although this region did not show IL-4 induced accessibility, it appeared to be preferentially open in non-cycling MF clusters (C5 and C1). In addition, we detected IL-4-induced RNAPII occupancy at this element. Measurement of eRNA expression identified G1-biased activity in the M0 state, and IL-4 exposure robustly induced eRNA production in a strongly G1-biased manner **(Figure 4G)**. These results identify the putative enhancer elements that likely drive the observed CC phase-biased expression of *Mgl2* and *Retnla*.

### IL-4 priming imprints a memory chromatin signature in a subpopulation of MFs and is limited by cell cycle

Our results show that MFs enter different plasticity states in a CC-phase-dependent manner. Therefore, we wondered if CC might also affect other aspects of MF responses, such as memory formation at the chromatin level. We used IL-4 for these experiments, since this cytokine has been shown to reprogram MF responses to secondary stimuli, indicative of a stable chromatin imprint [22, 23, 28]. Hence, we established a MF priming model in which IL-4 polarization (24h) is followed by cytokine washout and resting (24h – IL-4-primed; referred to as M2p) and performed scATAC-seq **(Figure 5A)**. Dimensionality reduction followed by UMAP of the M0, M2, and M2p MFs (n=18,376) suggested that the chromatin structure of M2 MFs is not stable after the removal of IL-4. Notably, the majority (~95%) of the cells from the M2p condition did not colocalize with either the M2, or the M0 states, suggesting a largely transient IL-4-induced chromatin imprint but also indicating a unique chromatin structure of primed MFs **(Figure 5B)**.

**Figure 5.**
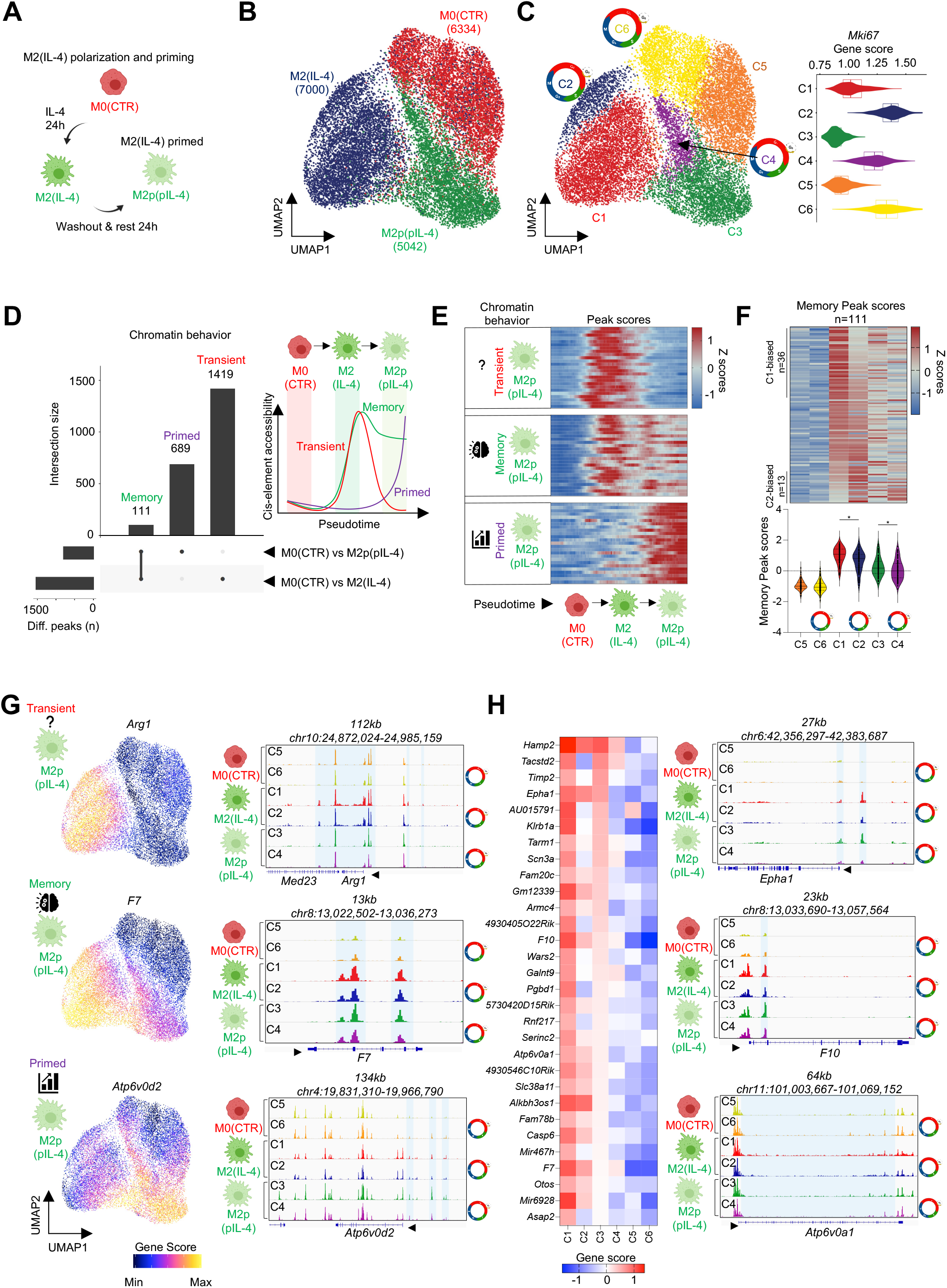
Cell cycle negatively affects the formation of memory in a subset of macrophages at the chromatin level. **(A)** Scheme on the model of M2 polarization and priming (M2p). **(B)** UMAP of M0(CTR), M2(IL-4) and the primed M2p(pIL-4) macrophage states. **(C)** UMAP colored by the 6 clusters identified. Violin plot depicts the gene score of *Mki67* in the 6 clusters. Cell cycle icons highlight clusters of cells predicted to be in cell cycle. **(D)** Upset plot of the differentially accessible cis-elements in M0(CTR) vs. M2p(pIL-4) and M0(CTR) vs. M2(IL-4) comparisons, and their overlap, yielding “memory”, “primed” and “transient” chromatin features (cis-regulatory elements). Scheme represents the behavior of these chromatin features across the conditions. **(E)** Heatmap of peak scores exhibiting distinct chromatin remodeling dynamics over the pseudotime of M2 polarization and priming. 25 peaks are shown **(F)** Heatmap visualization of the “Memory” peak scores from panel D across the 6 clusters. Peaks with significantly biased accessibility scores are highlighted in C1 and C2. Violin plot representation of the distribution of peak scores across the clusters (bottom). Wilcoxon Signed Rank Test, p<0.0001. **(G)** UMAP visualization of the gene score values for the indicated genes that display different chromatin remodeling activities in the 3 conditions (left). Genome browser views of scATAC-seq signal from the 6 clusters on the indicated gene loci (right). **(H)** Heatmap of cluster 1 marker gene scores (top30) over all clusters (left). Genome browser views of scATAC-seq signal from the 6 clusters on the indicated gene loci (right). Cell cycle icons highlight clusters of cells predicted to be in cell cycle.

Clustering MFs based on their chromatin states resulted in 6 clusters (M0 – C5 and C6; M2 – C1 and C2; M2p – C3 and C4). Among these, C2, C4 and C6 showed high *Mki67* accessibility, which reliably identifies cycling cells (Figure 3A), thus we designated them as CC-clusters **(Figure 5C)**. Identification of the specific CREs of the clusters revealed that the majority of IL-4-induced chromatin changes are lost after the removal of the cytokine, but cells in the primed state displayed a specific CRE program (C3 and C4) (FDR ≤0.01, Log_2_ FC≥1) **(Figure S5B)**. Notably, we found 1,641 IL-4-induced CREs, of which 1,530 returned to the steady state (“Transient”) and 111 showed persistent accessibility following IL-4-washout (“Memory”). We also observed 689 CREs that exhibited induced accessibility following IL-4-washout (“Primed“) **(Figure 5D, Table S7)**. As expected, these groups of CREs followed the anticipated accessibility patterns along the trajectory of priming by connecting and studying the transitional chromatin states of the 3 conditions (M0 – M2 – M2p) **(Figure 5E and S5A)**. Importantly, the accessibility of the “Transient” and “Primed” CREs were impacted in the CC-clusters. Specifically, 233 (C1-biased) and 21 (C2-biased) of the “Transient” CREs showed either reduced or increased accessibility in CC, respectively. Among the “Primed” CREs, 85 (C3-biased) and 8 (C4-biased) were either reduced on increased in CC, respectively **(Figure S5B)**. CREs with “Memory” characteristics followed a similar trend and showed 36 (C1-biased) and 13 (C2-biased) genomic regions with either reduced or increased accessibility in CC, respectively. We calculated the median peak score values of the “Memory” CREs in each cluster and found that both the establishment (comparing C1 to C2) and stability (comparing C3 to C4) of this chromatin imprint was negatively affected by CC (Wilcoxon Signed Rank Test, p<0.0001) **(Figure 5F)**. Finally, we annotated the CREs from the 3 groups to their putative target genes based on co-accessibility (see Methods) and proximity (200kb window around the gene TSS, **Table S7**). As expected, annotated genes also featured similar chromatin remodeling dynamics as the annotated CREs as judged by their gene score values (e.g., *Arg1* – “Transient”, *F7* – “Memory”, *Atp6v0d2* – “Primed”) **(Figure 5G)**.

Lastly, we identified genes exhibiting cluster-biased chromatin accessibility in the context of priming (FDR ≤0.01, Log_2_ FC≥1.25). We focused on the ones that displayed C1- (non-cycling) or C2-biased (CC-cluster) accessibility scores from the M2 polarized condition to identify IL-4-induced chromatin remodeling events. As a result, we found 102 genes with C1- and 85 genes with C2-biased accessibility **(Figure S5C)**. Visualization of the top 30 gene scores showed that C1-biased genes are strongly induced by IL-4 polarization and their accessibility is preferentially retained in C3 (non-cycling) of the primed cells **(Figure 5H)**. Furthermore, several of these genes had annotated memory CREs in their proximity (e.g., *Epha1*, *F10* and *Atp6v0a1*) **(Figure 5H)**. In contrast, the top 30 genes with C2-biased accessibility demonstrated strong CC-induced chromatin remodeling events, including CC-genes (e.g., *Top2a* and *Tubb5*), and IL-4 had effects on only 50% of the genes (e.g., *Clec10a* and *Rnase2a*) **(Figure S5D)**. Altogether, our results identify “Transient”, “Memory” and “Primed” CREs. CC negatively affects the establishment of the majority of CREs in these groups, including the memory chromatin imprint in a subset of IL-4-primed MFs. Therefore, CC limits chromatin remodeling events during MF priming with IL-4, providing an additional example of CC-influenced MF response.

### IL-4 priming and cell cycle limits repolarization by IFNG at the chromatin level

After we provided multiple lines of evidence that CC affects MF plasticity during M1/M2-polarization and priming, we set out to study an additional widely used system to test MF plasticity, called repolarization. In the experimental setting of repolarization, MFs are first polarized with either an M1 or an M2 cytokine followed by the treatment of the opposing polarization signal to assess the plasticity of the underlying polarized state [20, 23]. To study repolarization capacity, we performed IL-4-priming according as previously described **(Figure 5A)**, but after the resting period we exposed the cells to IFNG for 3 hours (repolarization) and performed scATAC- and scRNA-seq **(Figure 6A)**. Using the single cell chromatin maps, we projected our 4 conditions (M0, M2p, M1 and repolarized – M1rep(pIL-4+IFNG)) and observed that the M1rep cells clustered close to the M1 polarized cells but only slightly overlapped with this condition on the UMAP **(Figure 6B)**. We identified 2 clusters in each condition (8 clusters total) **(Figure 6C)**. Importantly, one of the 2 identified clusters in each condition were CC-clusters exhibiting strong accessibility for CC genes (C1 – M1; C4 – M1rep; C5 – M0 and C8 – M2p), such as *Top2a*, *Ccnf*, *Hist1h2bb* and *Mki67* **(Figure 6D and S6A)**, further reinforced by the expression (gene integration score) of *Mki67* **(Figure S6B)**.

**Figure 6.**
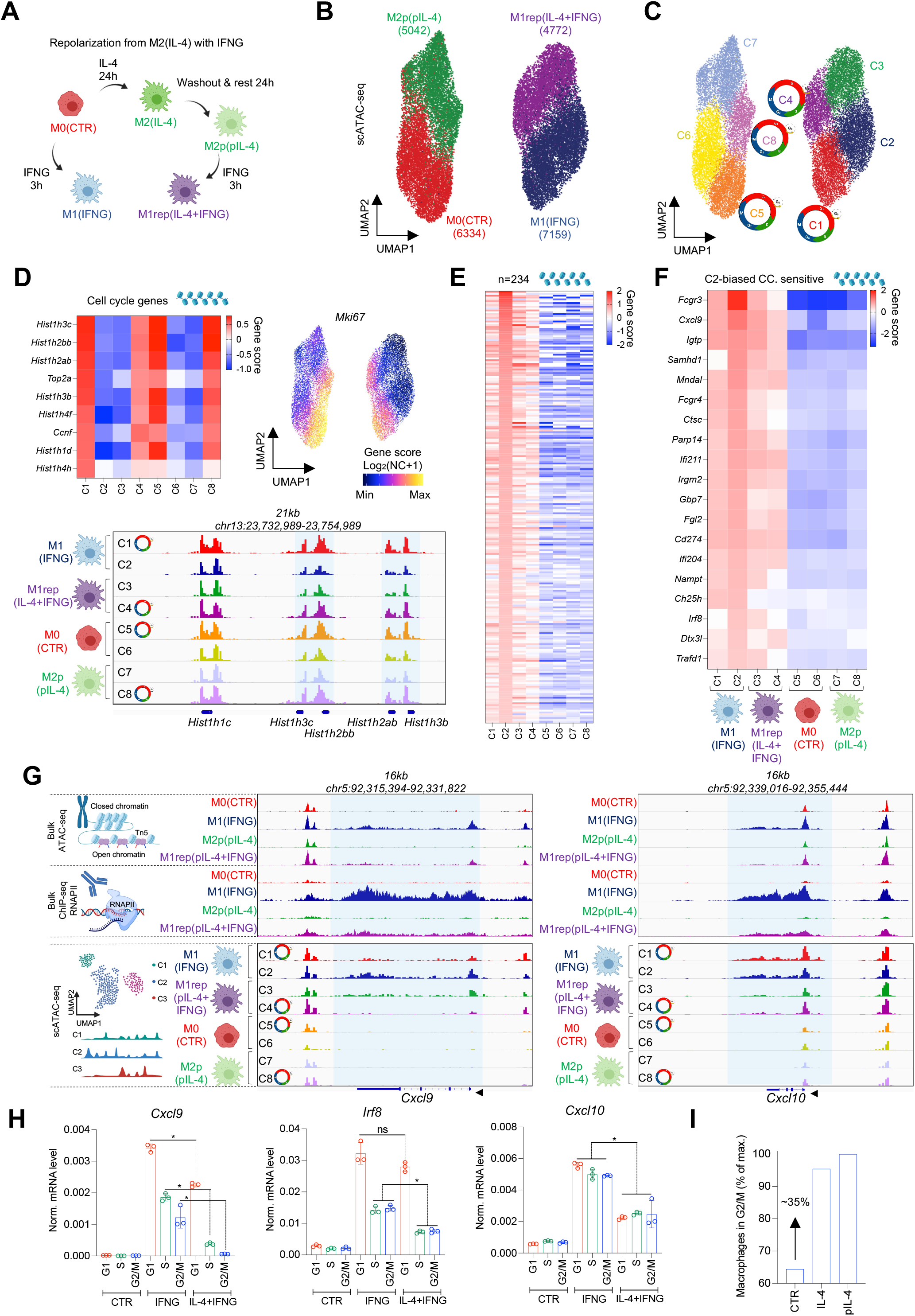
IL-4 priming cooperates with cell cycle to limit the repolarization capacity of IFNG in a subset of macrophages. **(A)** Scheme on the model of IL-4 priming and repolarization by IFNG. **(B)** UMAP of M0(CTR), M2p(pIL-4) – IL-4-primed, M1(IFNG) and M2repol(pIL-4+IFNG) – IL-4-primed and repolarized with IFNG conditions. **(C)** UMAP colored by the 8 chromatin clusters. Clusters of cells predicted to be in cell cycle are indicated with cell cycle icons. **(D)** Heatmap of cell cycle marker gene scores. UMAP of *Mki67* gene score values. Log_2_ normalized counts+1 is shown. Genome browser snapshot of scATAC-seq signal on cell cycle genes in the 8 clusters. Clusters of cells predicted to be in cell cycle are indicated with cell cycle icons. **(E)** Gene score heatmap of cluster 2 markers across all clusters. **(F)** Gene score heatmap of the markers of cluster 2, also detected as IFNG-induced, G1-phase-biased transcripts by bulk RNA-seq on Figure S4D. **(G)** Genome browser views on *Cxcl9* and *Cxcl10*. Bulk ATAC- and RNAPII ChIP-seq signals in CTR, IFNG, IL-4-primed and IL-4-primed and repolarized conditions (top part). scATAC-seq signal in the 8 clusters (bottom part). Clusters of cells predicted to be in cell cycle are indicated with cell cycle icons. **(H)** mRNA levels of *Irf8*, *Cxcl9* and *Cxcl10* in cell cycle across M0(CTR), M1(IFNG) and M2rp(pIL-4+IFNG) conditions. On the bar graphs, significant changes were determined by two tailed, unpaired t-test at p<0.05 (n=3). Shown are means with SDs. **(I)** Percentage of macrophages in the G2/M phase of the cell cycle as determined by FACS. Average of 3 experiments are used to calculate the percentage-wise distribution of cells in G2/M relative to the highest value (M2p).

Next, we defined the IFNG-induced chromatin remodeling events that are free from the effects of CC, using non-cycling M1 MFs from C2 **(Figure 6C, D)**. We identified 234 marker genes in C2 and found that these genes exhibited significantly weaker accessibility in the M1 MF CC-cluster (C1) (FDR ≤0.01, Log_2_ FC≥1), reinforcing our RNA-seq results **(Table S8)**. Interestingly, we also detected reduced accessibility of these genes in non-cycling M1rep MFs in C3, suggesting that IL-4-priming has a similar effect on the chromatin state of these genes as CC. Furthermore, MFs in C4 (M1rep MFs in the CC-cluster) displayed an even more severe defect in chromatin remodeling upon IFNG stimulation, suggesting that CC and IL-4-priming shift this subpopulation into an “IFNG-tolerant” plasticity state **(Figure 6C and E)**. To link these findings with the RNA-seq results of CC-phase sorted MFs, we overlapped the IFNG-induced, G1-biased gene signature (Figure S4D; 41 genes) with the marker gene scores of C2. This yielded a list of genes that are negatively affected by CC at both the gene expression and chromatin level, while IL-4 priming also reduced their sensitivity to IFNG at the chromatin level (n=19; e.g., *Fcgr3*, *Cxcl9*, *Fcgr4*, and *Irf8*) **(Figure 6F)**. These results indicate that IL-4-priming and CC might work in an additive or even in a synergistic fashion to further reduce MF plasticity to IFNG at the subpopulation level.

We tested this hypothesis on three IFNG-induced genes, two of which are CC-sensitive, *Cxcl9* and *Irf8*; and one that is CC-insensitive, *Cxcl10* (as revealed by RNA-seq). First, we performed RNAPII ChIP-seq in the repolarization system. Second, we visualized aggregated scATAC-seq with the RNAPII ChIP-seq signal from M0, M1, M2p and M1rep conditions on the loci of *Cxcl9* and *Cxcl10* **(Figure 6G)**. As expected, IFNG-induced chromatin remodeling and RNAPII recruitment was reduced in the M1rep condition compared to the M1 state. Importantly, the two gene loci appeared to behave essentially the same in the bulk datasets. However, when we visualized the scATAC-seq signal in each cluster, we observed striking differences. *Cxcl9* exhibited reduced accessibility in cycling, M1 polarized cells (C1) and also after IL-4 priming in non-cycling cells (C3). We detected an even lower level of accessibility in IL-4-primed, cycling MFs (C4) **(Figure 6G and S6A)**. In addition to this, we observed similar chromatin accessibility changes around *Irf8* **(Figure S6A)**. In contrast, *Cxcl10* did not show differences between cycling (C1) and non-cycling (C2) M1 polarized MFs. Moreover, we detected uniformly reduced chromatin accessibility in both cycling (C4) and non-cycling (C3) M1rep MFs **(Figure 6G)**. Gene integration scores from the scRNA-seq experiments also supported these observations on all of these genes **(Figure S6B)**.

In order to provide experimental evidence that CC and IL-4-priming can limit IFNG responsiveness at the subpopulation level, we measured gene expression from CC phase sorted MFs from M0, M1 and M1rep conditions. *Cxcl9* and *Irf8* exhibited reduced expression levels in the S and G2/M phases of CC upon IFNG treatment validating the bulk RNA-seq results. However, we detected greater reduction of the IFNG response in the S and G2/M phases of CC from M1rep MFs, confirming our previous results that CC and IL-4-priming are two major factors that can limit MF plasticity to IFNG. Moreover, CC and priming appeared to reduce IFNG response in a synergistic or additive fashion on *Cxcl9*, or *Irf8*, respectively. In contrast, *Cxcl10* was not sensitive to CC, and IL-4-priming uniformly reduced MF plasticity to IFNG in all CC-phases **(Figure 6H)**.

Since IL-4 has been described to induce MF proliferation *in vivo*, we wondered if the M2 and M2p MF populations display differences in their CC-phase distribution, which can provide an additional mechanism to limit IFNG responsiveness according to our previous results [38]. Using FACS, we quantified the CC-phase distribution of M2 and M2p MF populations and have not detected significant differences in the G1- or S-phase but observed a 35% increase in MF numbers in the G2/M-phase of CC **(Figure 6I and S6D)**. Therefore, our results provide strong evidence that MF plasticity to IFNG is reduced in CC, and IL-4 priming achieves very similar effects in non-cycling cells. Surprisingly, plasticity to IFNG has dramatically changed in cycling, M2p MFs, suggesting that priming with IL-4 and CC are two major and complimentary factors that can affect MF plasticity at the subpopulation level. Furthermore, IL-4 can change the CC-phase distribution of the population, directing more MFs into the G2/M-phase, representing an additional mechanism to limit IFNG response at the population level.

### MFs express a cell cycle-intrinsic tissue remodeling gene program

Our findings show that CC determines the plasticity of MFs to environmental changes (i.e., polarization signals) but whether there are CC-intrinsic gene expression programs which might support specialized macrophage functions is unknown. Therefore, we sought to study the gene expression program of CC-phase sorted (G1, S and G2/M) M0 macrophages by RNA-seq. Differential gene expression analysis of a three-way comparison across the CC-phases identified 3,776 CC-sensitive and 7,327 insensitive genes (Benjamini–Hochberg adjusted p-value ≤0.001; FC≥1.3) **(Figure 7A and S7A)**. We performed pathway analysis on the CC-sensitive genes, which showed CC-related functional categories, including “Kinetochore metaphase signaling pathway” and “Cell cycle control of chromosomal replication”. Surprisingly, we also noted functional terms that are related to fibrosis and tissue remodeling, for example, “Hepatic fibrosis/Tissue remodeling” **(Figure 7B)**. As expected, the first two terms mainly described the known gene expression program of the S (e.g., *Pcna*, *Mcm2* and *Pola1*) and G2/M (e.g., *Cenpp*, *Cenpe* and *Spdl1*) phases of CC **(Figure 7C)**. However, genes in the fibrosis and tissue remodeling term also exhibited S-G2/M-phase-biased expression (*Mmp9*, *Col1a1*, *Col1a2*, *Acta2*, *Fn1* and *Vcam1*). We sorted MFs and validated the CC phase-dependent induction of *Fn1*, *Acta2* and *Col1a1*, reproducing the RNA-seq results **(Figure 7D)**. Collectively, these results uncover a CC-intrinsic gene expression signature that is linked to tissue remodeling and fibrosis, preferentially expressed in the S-G2/M phases of CC. Interestingly, these genes are linked to tissue regeneration in different model systems of wound healing and muscle regeneration after injury, suggesting the potential relevance of cycling MFs during *in vivo* conditions [16, 28]. Additionally, MF proliferation has been noted as a feature of the resolving phases of inflammatory processes and regeneration, raising the question whether proliferating MFs can express the uncovered tissue remodeling genes under *in vivo* circumstances [14, 16, 17].

**Figure 7.**
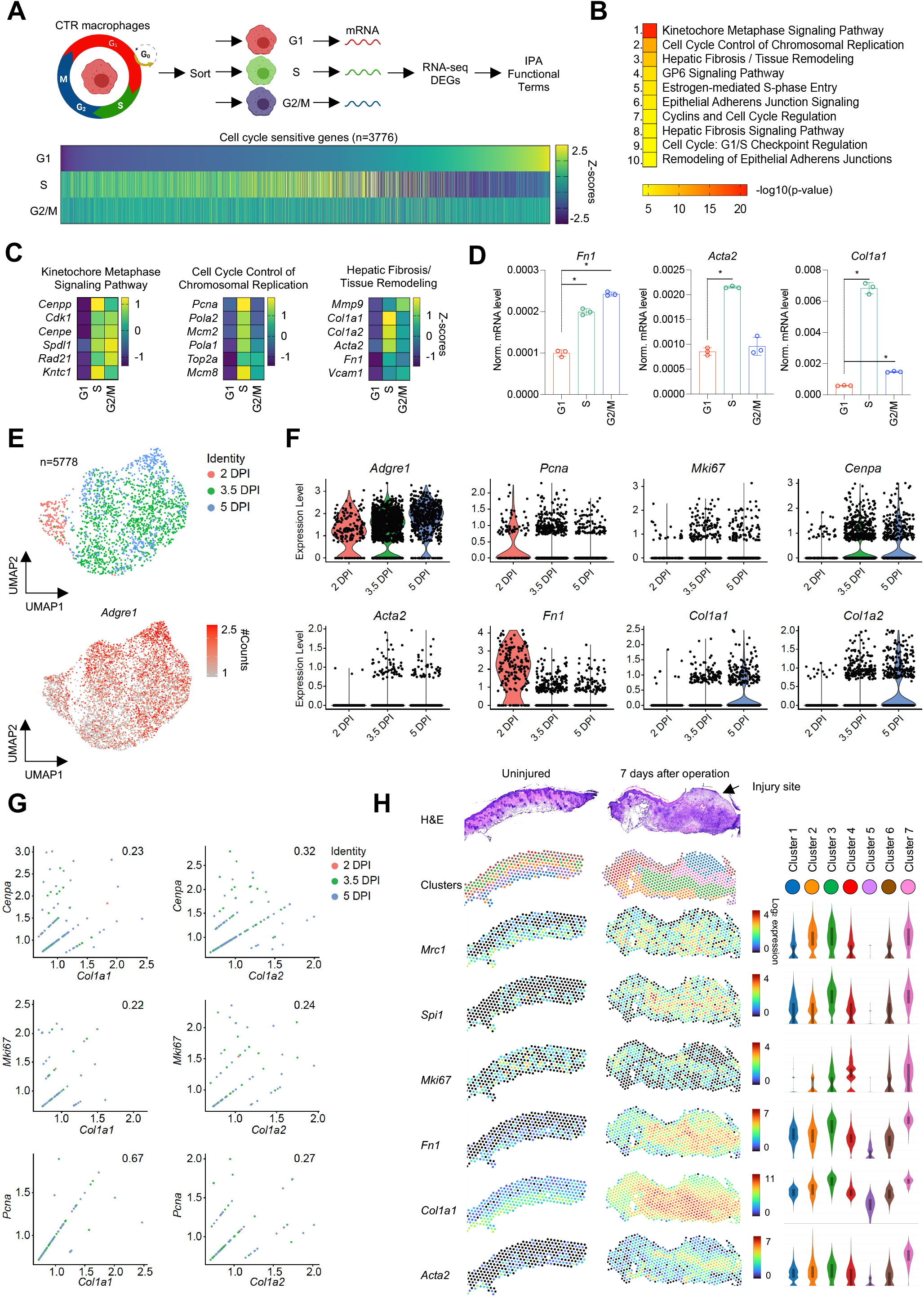
Cycling macrophages upregulate a tissue regeneration gene program. **(A)** Scheme of the experimental setting. DEGs – differentially expressed genes, IPA – Ingenuity Pathway Analysis. Heatmap represents differentially expressed genes across the cell cycle phases in M0(CTR) macrophages. **(B)** Ingenuity Pathway Analysis of the differentially expressed genes. Top 10 enriched biological functions are shown. **(C)** Expression of a select set of genes from the first three enriched biological functions are shown determined by bulk RNA-seq. **(D)** mRNA levels of *Fn1*, *Acta2* and *Col1a1* measured by RT-qPCR. Significant changes were determined by two tailed, unpaired t-test at p<0.05 (n=3). Shown are means with SDs. **(E)** UMAP projection of macrophages from regenerating muscle, 2-, 3.5- and 5-days post-injury (DPI) determined by scRNA-seq (GSE138826) (top). UMAP of the expression values of *Adgre1* (F4/80) (bottom). # - normalized. **(F)** Violin plots represent gene expression of the indicated genes in single macrophages on the different days (DPI) of regeneration (LogNormalized Expression). **(G)** Feature scatter plots of the indicated gene pairs visualizing co-expression in single macrophages (LogNormalized Expression). Single cells are colored by days post-injury (DPI). Pearson correlation coefficients are indicated on each plot. **(H)** Loupe browser images on spatial transcriptomics results in the stented wound healing model. Hematoxylin and eosin (H&E) staining of the tissue is shown on top. Clusters defined based on the expression programs of different tissue layers is shown below. Expression levels of individual genes are depicted on the tissue slides and on the violin plots in the 7 clusters (Log_2_ Expression values).

### Proliferating MFs express tissue remodeling factors during tissue regeneration

MFs are indispensable for proper muscle regeneration [44–46]. We reanalyzed single cell transcriptomic datasets of regenerating muscle, which exhibits monocyte - MF differentiation in the inflammatory phase and MF proliferation in the regenerative phase. Importantly, in the regenerative phase, MFs can support angiogenesis and extracellular matrix remodeling, which is required for regeneration [15, 16]. We processed a dataset of cardiotoxin (CTX)-induced muscle injury of the *tibialis anterior* muscle of mice, where regeneration was followed at 7 different time points until day 21 (days 0, 0.5, 2, 3.5, 5, 10 and 21) [47]. We subset MFs based on the expression of *Adgre1* (F4/80), *Mrc1*, *Msr1* and *Itgam* (LogNorm.Expression>0.5); and found 5,997 cells exhibiting the combinations of these features. We observed a massive surge in MF numbers starting at day 2 (n=537 versus day 0.5 n=15), that peaked at day 3.5 (n=3,655) and began to decline at day 5 (1,586). By days 10 (n=134) and 21 (n=68), MF count almost returned to the baseline, where damaged myofibers are regenerated (days post injury - DPI10) and the tissue is fully restored (DPI21), respectively **(Figure S7B)** [36]. Due to these temporal patterns in MF numbers, we subset MFs from days 2, 3.5 and 5 following the injury (n=5,778) and performed dimensionality reduction **(Figure 7E)**. MFs did not cluster based on CC, but rather, cycling cells appeared to be scattered in the low dimensional space between the different MF phenotypes based on CC-scoring and the expression of CC genes (*Pcna*, *Mki67* and *Cenpa*) **(Figure S7C)**. We detected more cells expressing these CC genes at days 3.5 and 5 compared to day 2 in line with our observations on the overall MF numbers observed at these days. Additionally, we found 4 of the 6 tissue regeneration genes also followed this expression pattern (*Acta2*, *Fn1*, *Col1a1* and *Col1a2*) **(Figure 7F)**. Next, we focused on MFs that express both the CC and the tissue remodeling genes (LogNorm.Expression>0.1); and performed a correlation analysis. We observed positive correlation between the expression of the collagen genes (*Col1a1* and *Col1a2*) and all three CC genes at the single cell level **(Figure 7G)**. Additionally, we found positive correlation between *Pcna* – *Acta2* and *Pcna* – *Fn1* expression supporting the finding that proliferating M0 MFs express this gene set. Although there was no correlation between either *Cenpa* - *Acta2* or *Cenpa* - *Fn1*, nor between *Mki67* - *Acta2* or *Mki67* - *Fn1* providing an internal negative control showing that the expression level of these genes in single MFs that express these genes are not always correlated **(Figure S7D)**.

Next, we analyzed another muscle regeneration dataset, in which regeneration was studied after barium chloride-induced *tibialis anterior* injury at days 4 and 7 after the challenge [48]. We subset MFs by the expression of *Mrc1*, *Msr1*, *Itgam* and *Adgre1* (LogNorm.Expression>0.5), performed dimensionality reduction and identified 5 clusters **(Figure S7E and S7F)**. In cluster 2, MFs expressed CC genes (*Mcm6*, *Pcna*, *Mki67* and *Cenpa*) along with tissue remodeling genes (*Fn1*, *Mmp9*, *Col1a1* and *Col1a2)* **(Figure S7G and S7H)**. We performed correlation analyses using MFs that co-express the CC markers and the collagen genes. Importantly, the collagen genes also showed positive correlation with all three CC-genes in this system as well **(Figure S7I)**.

Finally, we used a spatial transcriptomics dataset of a stented wound healing mouse model [49]. In this system, a stent is introduced into the dorsal skin of mice that more closely mimics human wound healing kinetics by disrupting the immediate wound construction mechanism achieved by the subdermal muscle layer called *panniculus carnosus*. After an incision is made, healing of the wound can be studied by collecting tissue sections for spatial transcriptomics **(Figure S7J)**. We utilized uninjured and injured (7 days post operation) spatial transcriptomic profiles generated by the 10x Genomics Visium platform. Clustering the tissue sections according to the transcriptional programs of each Visium spot yielded 7 clusters. Three of these clusters (1, 7 and 3) exhibited high expression of MF markers, which were virtually absent in the uninjured state (*Msr1*, *Mrc1* and *Spi1*) **(Figure 7H)**. We observed massive MF infiltration at the center of the injury site as noted previously, mainly corresponding to cluster 1 and 7 [38]. Interestingly, cluster 7 showed high expression for both *Mki67* and the MF marker genes. This same region in the dermis exhibited the highest expression of the tissue regeneration-related genes (*Fn1*, *Col1a1*, *Acta2*, *Col1a2*, *Mmp9* and *Vcam1*), where both MFs and fibroblasts can produce these gene products **(Figure 7H and S7F)**. In summary, our results reveal the expression of a CC-intrinsic tissue remodeling gene signature, which is preferentially expressed in the S-G2/M-phases of CC in M0 MFs. These muscle regeneration and wound healing models provide evidence that cycling MFs can express parts of this signature *in vivo*.

## Discussion

MFs exist in distinct plasticity states within tissues, and the overall heterogeneity of the population can be decisive when environmental factors perturb the homeostatic balance, for example in the presence of infection, injury, or cancer [50, 51]. Understanding the drivers of heterogeneity is of great interest due to the programmability of the MF niche, but the major determinants are still unknown. Here we provide evidence that one of the most fundamental biological processes, CC, influences MF plasticity.

MF proliferation is a general phenomenon across tissues during an immune challenge [17, 38, 52, 53]. Entering CC replenishes and maintains MF populations, although progression through the phases of CC can provide opportunities to support additional immunological functions, a concept that has not been covered. We coin the term, cyclical immune plasticity, which describes MF plasticity to polarization signals and immunological functions in the different phases of CC. We provide evidence for CC-impacted MF responses in three independent model systems: 1. polarization; 2. priming; and 3. repolarization.

In the MF polarization model, scATAC- and scRNA-seq uncovered heterogeneous responses to polarization cues, which coincided with CC markers. Single cell studies typically regress out CC effects that can strongly bias the clustering of immune cell populations. As a result, in these studies, CC-associated phenotypic traits remain hidden [54]. Here, we exploited this feature of our datasets to assess MF plasticity in CC. Importantly, we provide experimental evidence that MFs launch biased polarization programs in the different phases of CC, using two of the main polarization signals (IL-4 and IFNG). MF polarization by these and many other cytokines have been extensively studied, yet these studies have not implicated CC as a factor that can alter MF plasticity or immune functions [22, 23, 26, 27, 29, 55]. Strikingly, we report that the M1 polarization program is strongly restricted to the G1-phase of CC. In fact, IFNG and LPS triggered M1 polarization has been reported to arrest MFs in G1 or at the border of G1-S transition, perhaps to support the completion of the polarization process in agreement with our findings [37]. In contrast, although ~50% of the IL-4-induced gene expression program is also G1-biased, a significant part of M2 polarization occurs in a S-G2/M-biased fashion, including *bona fide* polarization marker genes, such as *Chil3*, *Mgl2* and *Fn1*. Moreover, we provide evidence that these gene expression patterns are carried out by phase-biased enhancer activities. Considering that IL-4 can induce MF proliferation *in vivo*, we propose that CC entry in conjunction with the IL-4-induced, phase-biased transcriptional programs might synergize to achieve heterogeneous polarization states that are collectively known as the “M2 polarized” phenotype at the population level [12, 13, 38].

MFs are notoriously one of the most plastic immune cell types [1, 2]. We tested this feature in an IL-4 priming model by performing polarization, cytokine washout, and rest. This system allowed us to study the transient and stable chromatin states of MFs. Population-level analysis of the chromatin states of MF polarization revealed mostly transient chromatin remodeling events after the removal of polarization signals, although stability has been also noted [22, 23, 28, 43]. Our single cell chromatin accessibility map in the priming model recapitulated the transient nature of the polarization process but identified a subset of “memory” MFs. This MF subset retained an IL-4-induced chromatin imprint around a specific set of polarization marker genes after cytokine removal. Importantly, we show that the memory imprint is sensitive to CC, thus cycling MFs cannot efficiently establish this chromatin state.

Several studies have employed opposing polarization signals and repolarization from a polarized state to mimic MF responses in complex immunological microenvironments [23, 28, 29]. These studies used bulk epigenome-mapping technologies and explained differences in MF responses solely by epigenetic effects that were established by the first stimuli, without providing single cell insights. Our repolarization model of M2 MFs with IFNG sheds light on a dampened inflammatory response, where CC and IL-4 priming work together to limit MF plasticity to IFNG at the subpopulation level. These findings provide further evidence for the roles of CC in shaping MF plasticity, and our results confirm that IL-4 priming can skew the CC-phase distribution of the population towards the G2/M-phase. Hence, MFs enter a highly restrictive plasticity state, not permissive to IFNG-induced transcription, which can also limit repolarization at the population level.

MF presence and proliferation is an apparent feature of regenerating tissues after injury or infections [14, 15, 16, 17]. Recent studies already noted the uncoupling of MF inflammatory and proliferative responses during infections of the lung and liver, connecting MF proliferation to the resolution of inflammation and regeneration [14, 17]. In agreement with this, our results indicate that MFs are less responsive to IFNG, while gain tissue remodeling gene expression programs in CC. Therefore, we propose that CC entry might provide a cyclical mechanism to dampen inflammation and support regeneration. Using published scRNA-seq datasets of regenerating muscle, we identified proliferating MF subsets, where the expression of tissue remodeling genes (*Col1a1* and *Col1a2*) and CC genes displayed positive correlation at the single cell level [47, 48]. Furthermore, spatial transcriptomics of wound healing also supported this concept, by defining a tissue layer of high MF-specific gene expression, along with the expression of the tissue remodeling gene set and *Mki67* [49]. These results imply that MFs not only change their plasticity states in the phases of CC but might also gain CC phase-intrinsic transcriptional programs, other than the ones that support DNA replication and cell division. Therefore, our findings are not only compatible with the growing recognition that MF proliferation aligns with the reparative phase of tissue injury and resolution of inflammation, but also puts forward the idea that MFs might obtain tissue regeneration-linked genetic programs in CC [14, 16, 17].

In summary, our results allow us to formulate the concept of cyclical immune plasticity using a model system of MF polarization. We propose that additional cell types of the immune system might use CC entry not only to replenish cell populations but also to tune their phenotype and level of plasticity, increasing cellular heterogeneity and flexibility at the population level. Future single cell studies should investigate and consider the role of CC as an immune regulatory process during infections and cancer. Finally, anti-cancer therapeutic applications targeting CC (e.g., CDK4/6 inhibitors) should be re-evaluated with respect to the immune cell community of the tumor microenvironment to understand how CC inhibition affects immune cell function [45].

## Limitations of the study

Although our results provide a few specific cases of CC phase-biased gene/enhancer activities, a more in-depth mechanistic understanding of CC-phase-biased gene expression is required. Additionally, identifying the TFs that drive phase-biased expression will be important future work for both the polarization-induced and the S-G2/M-intrinsic tissue remodeling gene set. Another caveat is the lack of knowledge on the mechanism by which IL-4 priming, and CC obtain similar, negative effects on MF IFNG response. We speculate that a still ongoing M2- or CC-driven gene expression programs might dampen MF plasticity to IFNG stimulation by squelching the basic transcriptional machinery and lowering the cells’ energy supply. Finally, although we use three different tissue regeneration models to provide correlative evidence for the appearance of the tissue remodeling gene set in cycling MFs *in vivo*, additional experiments will need to directly assess MF CC and its importance in tissue regeneration.

## Acknowledgments

We thank the members of the Satpathy and Chang labs for stimulating discussions, and M. Amouzgar and the informatics team at the Parker Institute for Cancer Immunotherapy for assistance with data analysis. This work was supported by the National Institutes of Health (NIH) K08CA23188-01 (A.T.S.), U01CA260852 (A.T.S.), RM1-HG007735 (H.Y.C.), a Career Award for Medical Scientists from the Burroughs Wellcome Fund (A.T.S.), a Technology Impact Award from the Cancer Research Institute (A.T.S.), an ASH Scholar Award from the American Society of Hematology (A.T.S.), the Parker Institute for Cancer Immunotherapy (H.Y.C., and A.T.S.), and the Scleroderma Research Foundation (H.Y.C.). H.Y.C. is an investigator of the Howard Hughes Medical Institute. J.A.B was supported by a Stanford Graduate Fellowship and a National Science Foundation Graduate Research Fellowship under Grant No. DGE-1656518. The sequencing data was generated with instrumentation purchased with NIH funds: S10OD018220 and 1S10OD021763. NIH NCI Postdoctoral Individual National Research Service Award 1F32CA239312-01A1 (D.S.F), the Advanced Residency Training at Stanford (ARTS) program (D.S.F.). NIH 1R01GM116892 (M.T.L), NIH 1R01GM136659 (M.T.L).

## Author contributions

B.D., H.Y.C and A.T.S conceptualized the study. B.D., J.A.B., H.Y.C., and A.T.S. wrote and edited the manuscript and all authors reviewed and provided comments on the manuscript. B.D., K.S., and Y.Q. performed experiments. J.A.B., S.L.M., A.Y.C., J.R.W., and H.K. analyzed data. B.D., H.Y.C. and A.T.S. guided experiments and data analysis. D.S.F., M.J., J.R.W., and M.T.L. performed analyses and provided single cell and spatial transcriptomics datasets.

## Data availability

Sequencing data has been deposited to GEO under accession: GSE178526

Published data that has been used in this study: GSE138826, GSE84520.

## Tables

Table 1: Gene scores of the trajectory analysis of M2 macrophage polarization. Related to Figure 1.

Table 2: Motif deviation scores of the trajectory analysis of M2 macrophage polarization. Related to Figure 1.

Table 3: Gene scores of the trajectory analysis of M1 macrophage polarization. Related to Figure 1.

Table 4: Motif deviation scores of the trajectory analysis of M1 macrophage polarization. Related to Figure 1.

Table 5: Cluster-biased gene score values of polarized macrophages. Related to Figure 2.

Table 6: Marker genes of M1 and M2 macrophages determined by scRNA-seq. Related to Figure S2B.

Table 7: Peak scores with transient, memory and primed kinetics. Related to Figure 5.

Table 8: Cluster-biased gene score values of repolarized macrophages. Related to Figure 6.

Table 9: Z-scores of cell cycle sensitive genes in M1 and M2 macrophages. Related to Figure 4.

Table 10: Primer sequences used in this study.

## Methods

### Bone marrow-derived macrophage culture

Wild type, 2-3 months old female C57Bl6 mice were purchased from Jackson laboratories. Mice were sacrificed and bone marrow was isolated form the tibiae and femora of the animals. Red blood cell lysis was carried out and cells were plated in differentiation media containing 10% FBS, Dulbecco’s Modified Eagle’s Medium (DMEM) and 20ng/ml mouse M-CSF (Peprotech). On the third day of differentiation, media was replaced with fresh differentiation media. Cytokine treatments sorting procedures were carried out on the 6^th^ day of differentiation.

### Treatment conditions

Macrophages were treated with either IL-4 (20ng/ml) or IFNG (20ng/ml) (Peprotech). For polarization we used 24 hours of IL-4 polarization and 3 hours of IFNG polarization in differentiation media that contained 10% FBS, 1% penicillin/streptomycin and M-CSF (20ng/ml). IL-4 priming was performed as follows: macrophages were polarized with IL-4 (20ng/ml) for 24 hours. Cell were washed three times with serum-free DMEM, then differentiation media was replaced, and cells were rested for an additional 24 hours. Repolarization was performed at this point for 3 hours with IFNG (20ng/ml). Macrophage polarization for cell cycle experiments used 3 hours of polarization with either IL-4 or IFNG (both at a 20ng/ml concentration).

### Fluorescence-activated cell sorting (FACS)

Macrophages (~3 x 10^6^) were stained with anti-F4/80 (rat monoclonal FITC-conjugated, BioLegend) in a 1:200 dilution in FACS buffer for 20 minutes on ice. Cells were spun and resuspended in serum-free DMEM pre-heated to 37C with Vybrant DyeCycle (1:500) and incubated for 30 minutes at 37C followed by Propidium Iodide (PI) staining and sorting. PI negative F4/80 positive macrophages were sorted from all three cell cycle stages according to the Vybant DyeCycle signal.

### Real-time quantitative PCR for enhancer RNA and mRNA detection (qPCR)

RNA was isolated with Trizol reagent (Ambion). RNA was reverse transcribed with High-Capacity cDNA Reverse Transcription Kit (Applied Biosystems) according to the manufacturer’s instructions. Transcript quantification was performed by qPCR reactions using SYBR green master mix (BioRad). Transcript levels were normalized to *Ppia*. Primer sequences are available from Table S10.

### Chromatin immunoprecipitation sequencing (ChIP-seq)

ChIP-seq was performed as previously described with minor modifications [1]. Bone marrow-derived macrophages (3 x 10^6^) were double crosslinked by 50mM DSG (disuccinimidyl glutarate, #C1104 - ProteoChem) for 30 minutes followed by 10 minutes of 1% formaldehyde. Formaldehyde was quenched by the addition of glycine. Nuclei were isolated with ChIP lysis buffer (1% Triton x-100, 0.1% SDS, 150 mM NaCl, 1mM EDTA, and 20 mM Tris, pH 8.0). Nuclei were sheared with Covaris sonicator using the following setup: Fill level – 10, Duty Cycle – 5, PIP – 140, Cycles/Burst – 200, Time – 4 minutes). Sheared chromatin was immunoprecipitated with RNAPIIpS2 antibody (Abcam - ab5095). Antibody chromatin complexes were pulled down with Protein A magnetic beads and washed once in IP wash buffer I. (1% Triton, 0.1% SDS, 150 mM NaCl, 1 mM EDTA, 20 mM Tris, pH 8.0, and 0.1% NaDOC), twice in IP wash buffer II. (1% Triton, 0.1% SDS, 500 mM NaCl, 1 mM EDTA, 20 mM Tris, pH 8.0, and 0.1% NaDOC), once in IP wash buffer III. (0.25 M LiCl, 0.5% NP-40, 1mM EDTA, 20 mM Tris, pH 8.0, 0.5% NaDOC) and once in TE buffer (10 mM EDTA and 200 mM Tris, pH 8.0). DNA was eluted from the beads by vigorous shaking for 20 minutes in elution buffer (100mM NaHCO_3_, 1% SDS). DNA was decrosslinked overnight at 65C and purified with MinElute PCR purification kit (Qiagen). DNA was quantified by Qubit and 10 ng DNA was used for sequencing library construction with the Ovation Ultralow Library System V2 (Tecan) using 12 PCR cycles. Libraries were sequenced on an Illumina Hiseq 2500 using paired-end 75bp reads.

### Bulk ATAC-seq and ChIP-seq computational methods

Bulk epigenetics datasets were analyzed as described previously [2]. Briefly, reads were trimmed for quality and adapter sequences using fastp. Trimmed reads were aligned to the mm10 reference genome using hisat2. Aligned reads were deduplicated using picard. Peaks were called for each sample using MACS2. A fixed-width, reproducible union peak set for each group of samples (e.g., bulk ATAC-seq samples) was constructed by iteratively merging individual peak calls for each sample and removing overlapping peaks until a final, non-overlapping set of peaks was obtained. The union peak set was used to create a sample by peak matrix. ATAC-seq coverage tracks were obtained by exporting normalized bigwig files from R, normalized to reads in TSS, a gold-standard normalization method that controls for both sequencing depth and library quality [3].

### Bulk RNA-seq

Approximately 20ng total RNA was used for library preparation with Ovation Ultralow RNA-seq V2 (Tecan) from two biological replicates. Libraries were generated according to the manufacturer’s instructions. Approximately 50ng amplified cDNA was subjected to Ovation Ultralow V2 library generation and manufacturer’s instructions were followed. Libraries were size selected with E-Gel EX 2% agarose gels (Life Technologies) and purified by QIAquick Gel Extraction Kit (Qiagen). Libraries were sequenced on HiSeq 2500 instrument.

### RNA-seq analysis

Fastq files were pseudoaligned to a mm10 transcriptome index and the abundance of transcripts was quantified using Kallisto v0.43.1 with bias correction [4]. The transcript-level abundance estimates were imported and summarized using tximport v1.16.1, and differential expression was determined using the DESeq2 package v1.28.11 in Bioconductor v.3.11. A gene was considered cell cycle-sensitive if it was differentially expressed between any two cell cycle stages in the control condition or the condition of interest (IL-4, or IFNG respectively) with an absolute fold change of ≥1.3 and a Benjamini– Hochberg adjusted p-value ≤0.001. If a gene was not differentially expressed between any two cell cycle stages with an adjusted p-value ≤0.001, it was considered cell cycle-insensitive. The cell cycle stage bias of a gene was assigned to the cell cycle stage where the gene showed the largest absolute scaled variance-stabilizing transformed expression.

### scATAC-seq sample and library generation

Single cell ATAC-seq experiments were performed on the 10x Chromium platform as described earlier [5]. Briefly, after cytokine treatments, macrophages were subjected to nuclei isolation according to the protocol of the manufacturer. Nuclei were counted and ~20,000 were submitted for tagmentation. After tagmentation, nuclei were loaded for capture using the 10x Chromium controller. After Gel emulsion generation, linear amplification was performed, followed by DNA purification following the manufacturer’s protocol. The resulting DNA was used for library construction as described on the website of the manufacturer. Libraries were quantified by quantitative PCR and were sequenced on an Illumina Hiseq 2500 sequencer, using the following setup: 50bp read 1N, 8bp i7 index, 16bp i5 index and 50bp read 2N. In this reaction, 1N and 2N refers to the DNA insert sequencing, while i5 and i7 sequencing identifies the individual barcodes of single cells.

### Single-cell RNA-seq library preparation

Single-cell RNA-seq libraries were prepared using the 10X Single Cell Immune Profiling Solution Kit (v1 Chemistry), according to the manufacturer’s instructions. Briefly, FACS sorted cells were washed once with PBS + 0.04% BSA. Following reverse transcription and cell barcoding in droplets, emulsions were broken, and cDNA purified using Dynabeads MyOne SILANE followed by PCR amplification (98°C for 45 sec; 14 cycles of 98°C for 20 sec, 67°C for 30 sec, 72°C for 1 min; 72°C for 1 min). For gene expression library construction, 50 ng of amplified cDNA was fragmented, and end-repaired, double-sided size selected with SPRIselect beads, PCR amplified with sample indexing primers (98°C for 45 sec; 14 cycles of 98°C for 20 sec, 54°C for 30 sec, 72°C for 20 sec; 72°C for 1 min), and double-sided size selected with SPRIselect beads. Single-cell RNA libraries were sequenced on an Illumina HiSeq 4000 to a minimum sequencing depth of 25,000 reads/cell using the read lengths 28bp Read1, 8bp i7 Index, 91bp Read2.

### scATAC-seq computational methods

scATAC-seq datasets were processed as described previously [6]. Briefly, reads were filtered, trimmed, and aligned to the mm10 reference genome using the 10X cellranger atac-count pipeline. Fragments files were loaded into ArchR for additional processing and analysis [7]. Separate ArchR projects were created for the three sample sets (priming, polarization, and repolarization) and additionally for each individual sample. Doublets were identified and removed using ArchR’s default doublet simulation and calling procedures. Barcodes were removed that had an enrichment of Tn5 insertions in transcription start sites (TSS enrichment) less than 4 or less than 1000 fragments. Tiles and GeneScores matrices were computed by summing Tn5 insertions in predefined genomic windows. After clustering the cells, peaks were called by macs2 on pseudoreplicates sampled from each cluster to obtain a reproducible peak set retaining cell type specific peaks. Transcription factor motif deviations were computed using chromVar [8]. Imputation was performed using Magic [9]. Pseudo-bulk tracks for indicated groups of cells were exported from ArchR as bigwig files normalized by reads in transcription start sites. Tracks were visualized in the Integrative Genomics Viewer (IGV).

### scRNA-seq computational methods

Reads were filtered, trimmed, and aligned to the mm10 reference genome using the 10X cellranger count pipeline. Doublets were called for each sample individually using the R implementation of scrublet [10], rscrublet. Gene by barcode counts matrices were loaded into Seurat for additional processing and analysis [11]]. Separate Seurat objects were created for the three sample sets (priming, polarization, and repolarization) and for each individual sample. Barcodes with >12.5% mitochondrial reads, <200 unique features, or a scrublet score >0.25 were removed. Remaining cells were then clustered and visualized. Cell cycle phase predictions for each cell were performed following the vignette available online: https://satijalab.org/seurat/archive/v3.1/cell_cycle_vignette.html. Published datasets were also analyzed according to these standards.

### Statistical methods

Statistical analyses were performed in R or GraphPad Prism. qPCR measurements were presented as means +/− SD and three biological replicates were performed. The exact replicate numbers are indicated in the figure legends for each experiment. On the bar graphs, significant changes were determined by two tailed, unpaired t-test at p<0.05. Differential chromatin accessibility analyses across cell clusters were performed with the following parameters: FDR ≤0.01, Log_2_ FC≥1.25, unless specified otherwise. Differential gene expression analyses of scRNA-seq results were performed with the following parameters: FDR≤0.01, FC≥1.3. Cell cycle phase-biased gene expression levels were determined as follows: Benjamini–Hochberg adjusted p-value ≤0.001; FC≥1.3 (two biological replicates were performed). Significant changes between the median peak scores of “Transient”, “Memory” and “Primed” chromatin regions were determined by Wilcoxon Signed Rank Test, p<0.0001. Statistical parameters are reported in the figure legends and also in the results section.

## Data availability

Sequencing data has been deposited to GEO under accession: GSE178526

Token for accessing the data: qrmtekckjjuxngd

Published data that has been used in this study: GSE138826, GSE84520.

