## Supplementary Text for "Macrophage inflammatory and regenerative response periodicity is programmed by cell cycle and chromatin state"

### Supplemental Information Legends

**Figure S1. Single cell chromatin accessibility landscape of MF polarization. Related to Figure 1.**

**(A)** Density plot of transcription start site (TSS) read enrichment as a function of unique scATAC fragment count per cell for the three samples. Cells passing the filter of having at least a TSS enrichment score of 3 and 1000 unique fragments are used in downstream analyses. Median TSS enrichment (MTE) is shown for each sample. **(B)** UMAP of TSS enrichment scores. **(C)** Tn5 bias corrected transcription factor footprints in M0(CTR), M2(IL-4) and M1(IFNG) macrophages around the respective motif's center. **(D)** Relative expression level of *Tlr2*, *Arg1* and *Itgax* from an IL-4 time course bulk RNA-seq experiment (GSE106706). **(F)** Heatmap visualization of the motif deviation scores of the indicated transcription factors over the M0(CTR) – M2(IL-4) and M0(CTR) – M1(IFNG) polarization trajectories. **(E)** scATAC-seq gene score values over the M2 polarization trajectory and relative bulk RNA-seq expression values (GSE106706) over the IL-4 time course for the indicated genes are shown, exhibiting: 1; reduced accessibility/expression (LOST), 2; gaining early accessibility/induction (EARLY) and showing late accessibility and late induction at the mRNA level (LATE). Small schematics indicate scATAC-seq data (Accessibility) and bulk RNA-seq data (mRNA level).

**Figure S2. Single cell transcriptomic analysis of MF polarization. Related to Figure 2.**

**(A)** UMAP of scRNA-seq experiments on M0(CTR), M2(IL-4) and M1(IFNG) macrophages. **(B)** Heatmap of differentially expressed genes in M0(CTR) vs. M2(IL-4) and M0(CTR) vs. M1(IFNG). Top20 induced and repressed genes are shown for both comparisons. Fold change over CTR is visualized. **(C)** Heatmap of integrated scRNA-seq expression values (Gene integration score), exhibiting biased expression level in the M2(IL-4) and M1(IFNG) scATAC-seq clusters. Red asterisks mark cell cycle genes. **(D)** ChromVAR transcription factor deviation scores (Dev.score) visualized either on the UMAP embeddings or as a ridge plot in the 6 clusters.

**Figure S3. Prediction of cell cycle state in polarized MFs indicate reduced polarization ability in cell cycle. Related to Figure 3.**

**(A)** Stacked bar plot depicts the distribution of macrophages across the 6 clusters as a function of their predicted cell cycle phase. **(B)** Predictions of cell cycle phase-biased gene expression using the integrated scRNA-seq values (gene integration matrix) projected onto the scATAC-seq clusters on panel Figure 3A. Comparisons across the different cell cycle phases are shown for both M2(IL-4) and M1(IFNG) exposed macrophages. **(C)** Gene scores and integrated gene expression values of *Egr2* (**(D)** for *Irf8*) are visualized on the UMAPs. Violin plots represent these values in the predicted cell cycle phases in either M2 (C) or M1 (D) polarized macrophages. **(E)** FACS gating strategy used to sort macrophages from distinct cell cycle phases.

**Figure S4. The transcriptional program of M1 and M2 polarized MFs in the different phases of cell cycle. Related to Figure 4.**

**(A)** Volcano plot of top50 induced and repressed genes upon IFNG polarization as determined by scRNA-seq. **(B)** Pie charts depict the percentage of cell cycle sensitive gene expression in the two polarization models (top). Smaller pie charts show the percentage of cell cycle sensitive induced or repressed genes in the two polarization models. **(C)** Genome browser views of IFNG-induced genes with G1-biased expression. **(D)** Heatmap of IFNG-induced genes exhibiting cell cycle phase-biased expression. **(E)** Heatmaps of IFNG- and IL-4-repressed genes with cell cycle sensitive expression. **(F, G)** Genome browser snapshots of genes with cell cycle-insensitive expression patterns in M1and M2 macrophages, respectively.

**Figure S5. The chromatin state of IL-4-mediated MF priming. Related to Figure 5.**
**(A)** UMAP of IL-4 priming trajectory (M0 – M2 – M2p). **(B)** Heatmap of the cluster specific cis-regulatory elements. Peak scores are visualized. Peaks with significantly biased accessibility are highlighted for the M2 and M2p conditions. **(C)** Gene score heat map of cluster specific marker genes. Gene scores with significant cluster-biased appearance are highlighted for the M2 and M2p conditions. **(D)** Gene score heatmap of cluster 2 markers (top30).

**Figure S6. MF repolarization is negatively affected by cell cycle. Related to Figure 6.**
**(A)** UMAPs and violin plots depict the gene score values (scATAC-seq) of the indicated genes. $Log_2$ normalized counts+1 is shown. **(B)** UMAPs and violin plots show the gene integration score values for the indicated genes. # - normalized **(C)** Representative FACS plot on the cell cycle phase distribution of M0, M2 and M2p macrophages (left). Percentage-wise cell cycle phase distribution of the samples (middle). Fraction of macrophages in the G2/M phase of cell cycle. Significant differences were identified by two tailed, unpaired t-test at $p<0.05$ (n=3) (right).

**Figure S7. Proliferating MFs of regenerating tissues express tissue remodeling genes. Related to Figure 7.**
**(A)** Scheme indicates the number of genes exhibiting cell cycle sensitive and insensitive expression patterns in M0(CTR) macrophages. Heatmap depicts cell cycle insensitive genes. **(B)** Macrophage numbers over the time course of regeneration in a cardiotoxin-induced injury model in the tibialis anterior muscle of mice (GSE138826). **(C)** scRNA-seq UMAP of macrophages colored by the clusters (left) or by cell cycle phase (right) (GSE138826). **(D)** Feature scatter plots of single macrophages that express the indicated genes. Pearson correlation coefficients are indicated on each plot (GSE138826). **(E)** scRNA-seq UMAP of macrophages derived from barium chloride injured tibialis anterior colored by the identified clusters. **(F)** Violin plots show the expression of macrophage markers, **(G)** cell cycle markers and **(H)** tissue remodeling associated genes. **(I)** Feature scatter plots of macrophages that express the indicated genes colored by the identified clusters on panel E from the barium chloride injury model. Pearson correlation coefficients are indicated on each plot. **(J)** Scheme of the stented wound healing model and the applied 10x Visium spatial transcriptomics platform. Loupe browser images on the tissue slides in uninjured and injured (7 days after injury) samples. $Log_2$ expression values are plotted on the tissue slides and in the identified clusters (violin plots).
