## Supplementary Figures for "Macrophage inflammatory and regenerative response periodicity is programmed by cell cycle and chromatin state"

### Supplementary Figure 1.

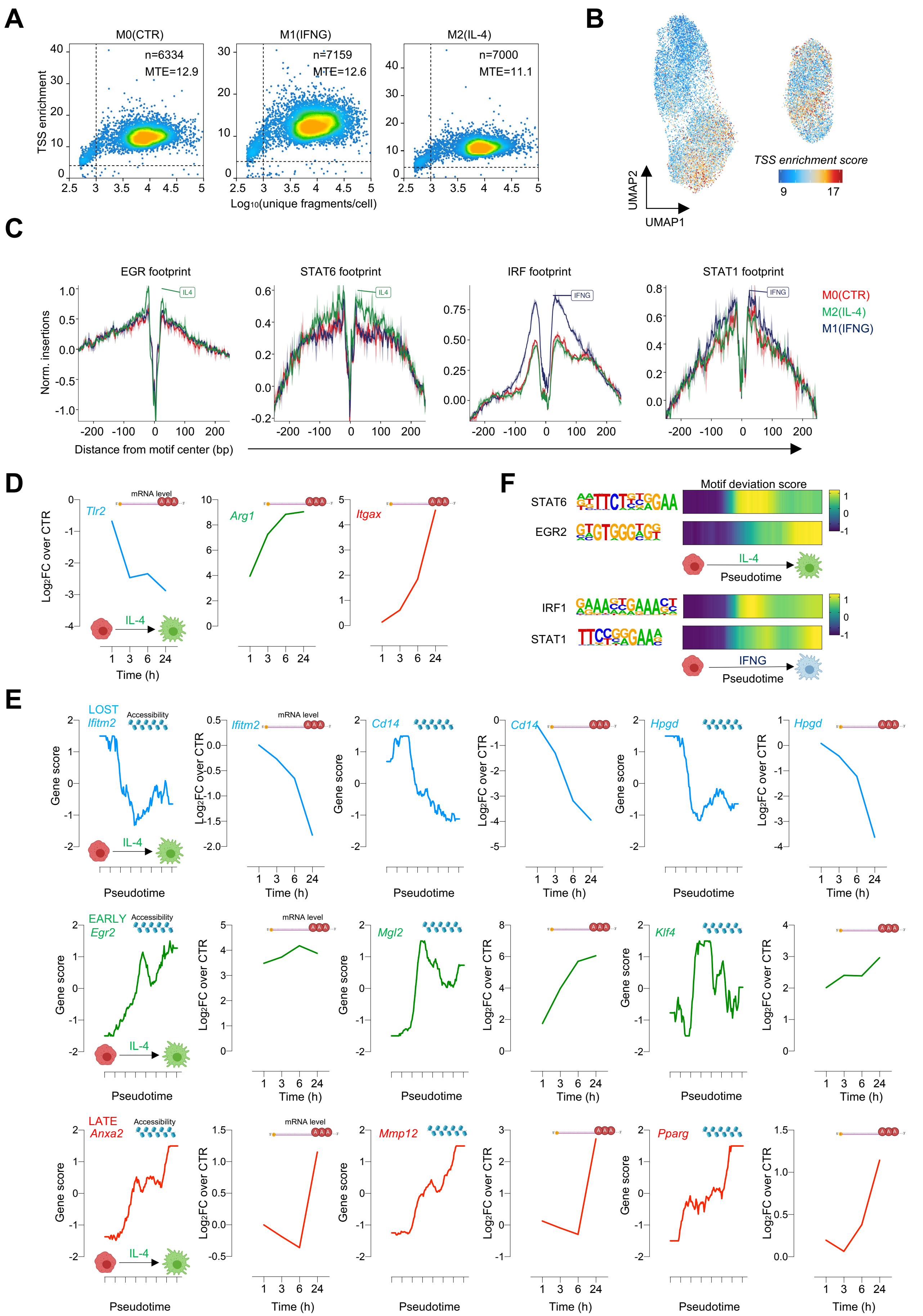

#### Supplementary Figure 2.

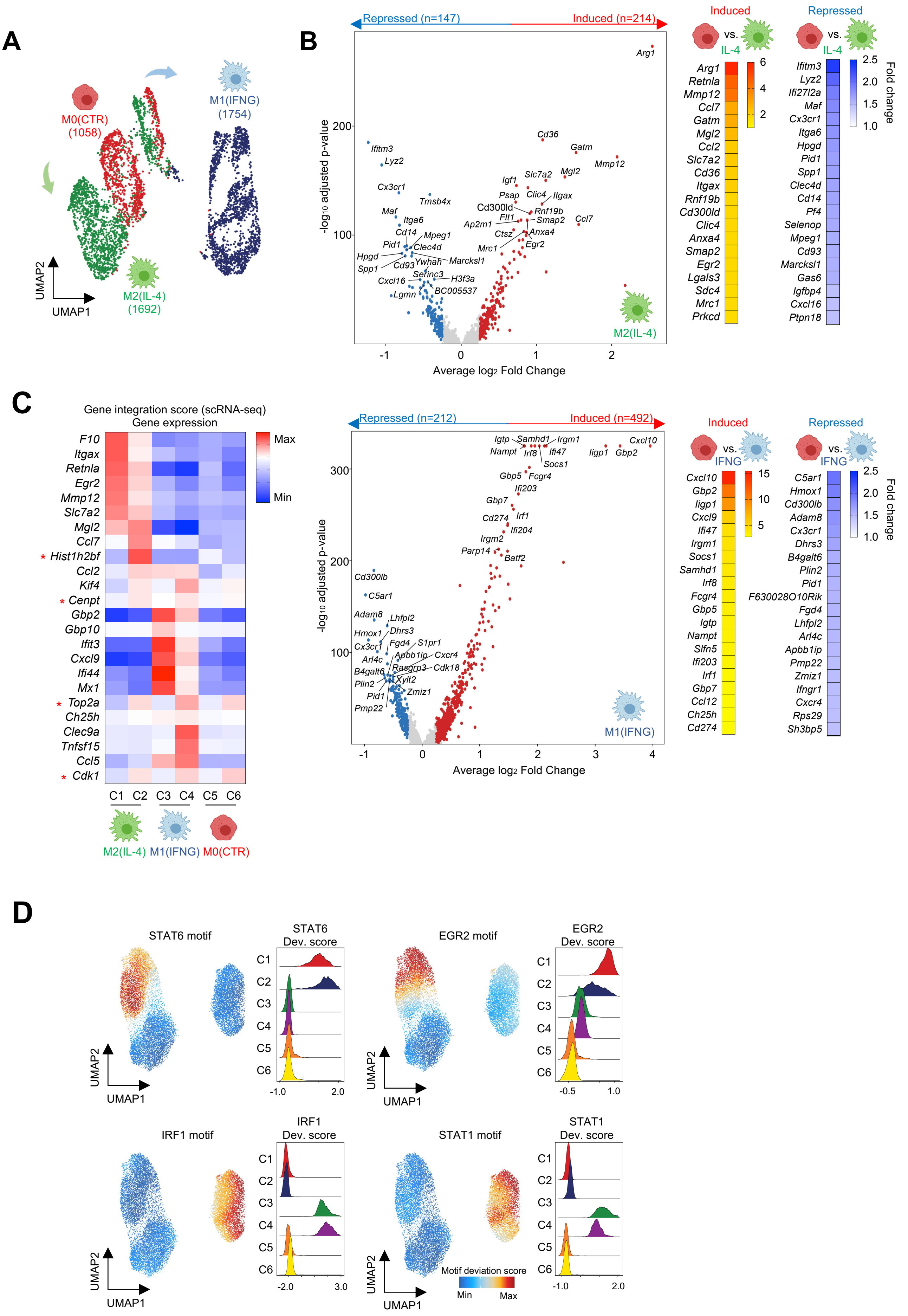

### Supplementary Figure 3.

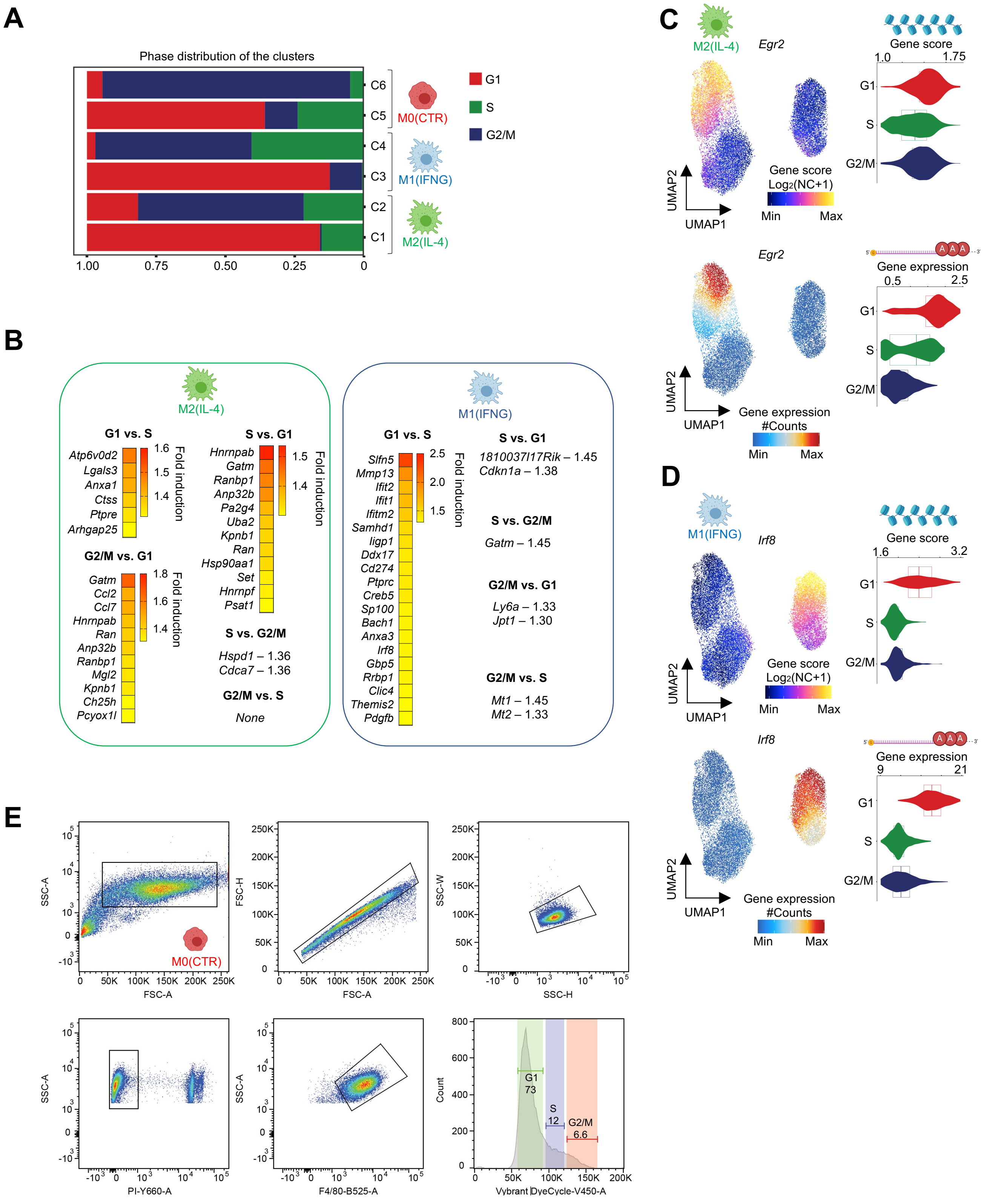

### Supplementary Figure 4.

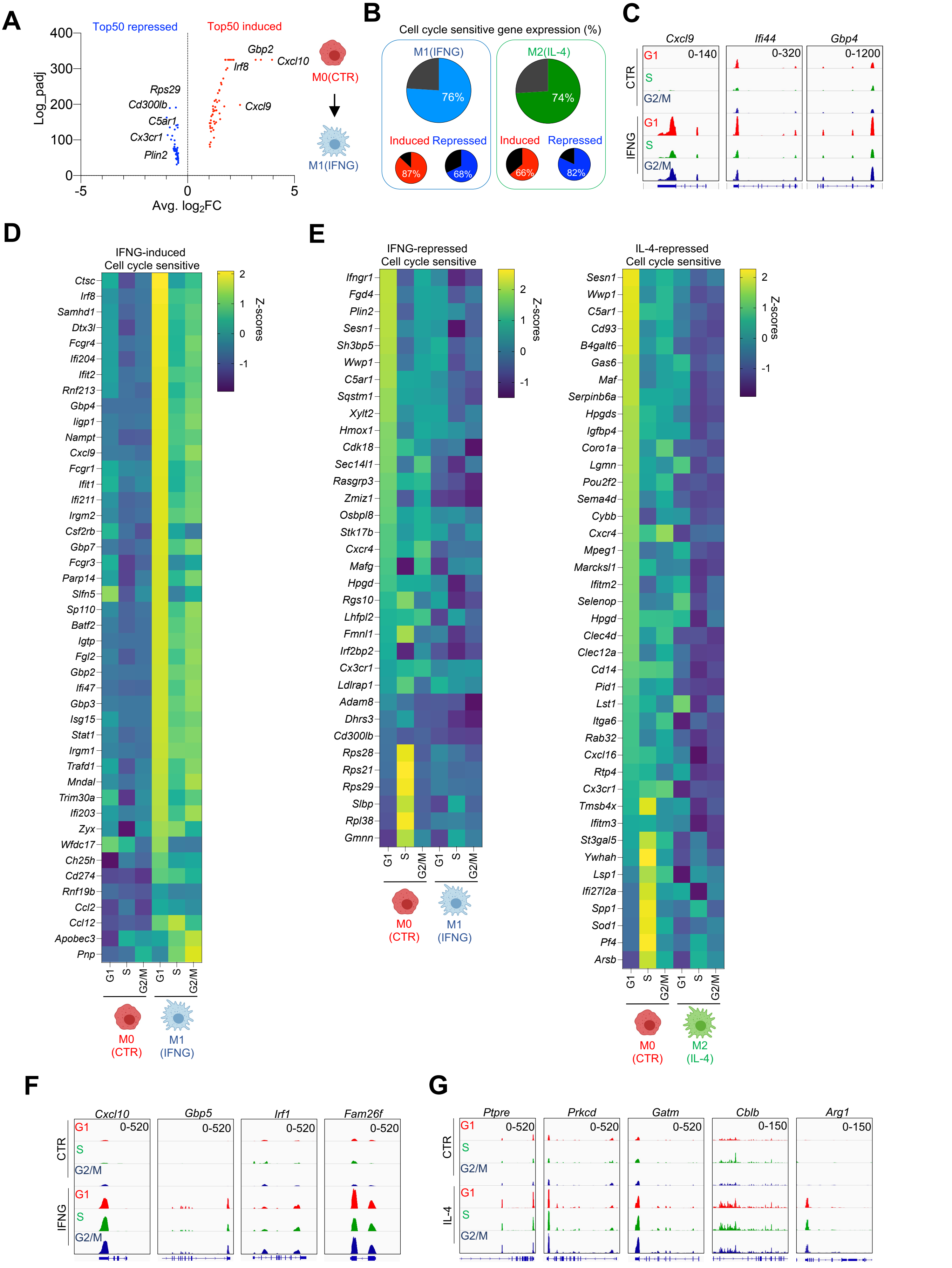

### Supplementary Figure 5.

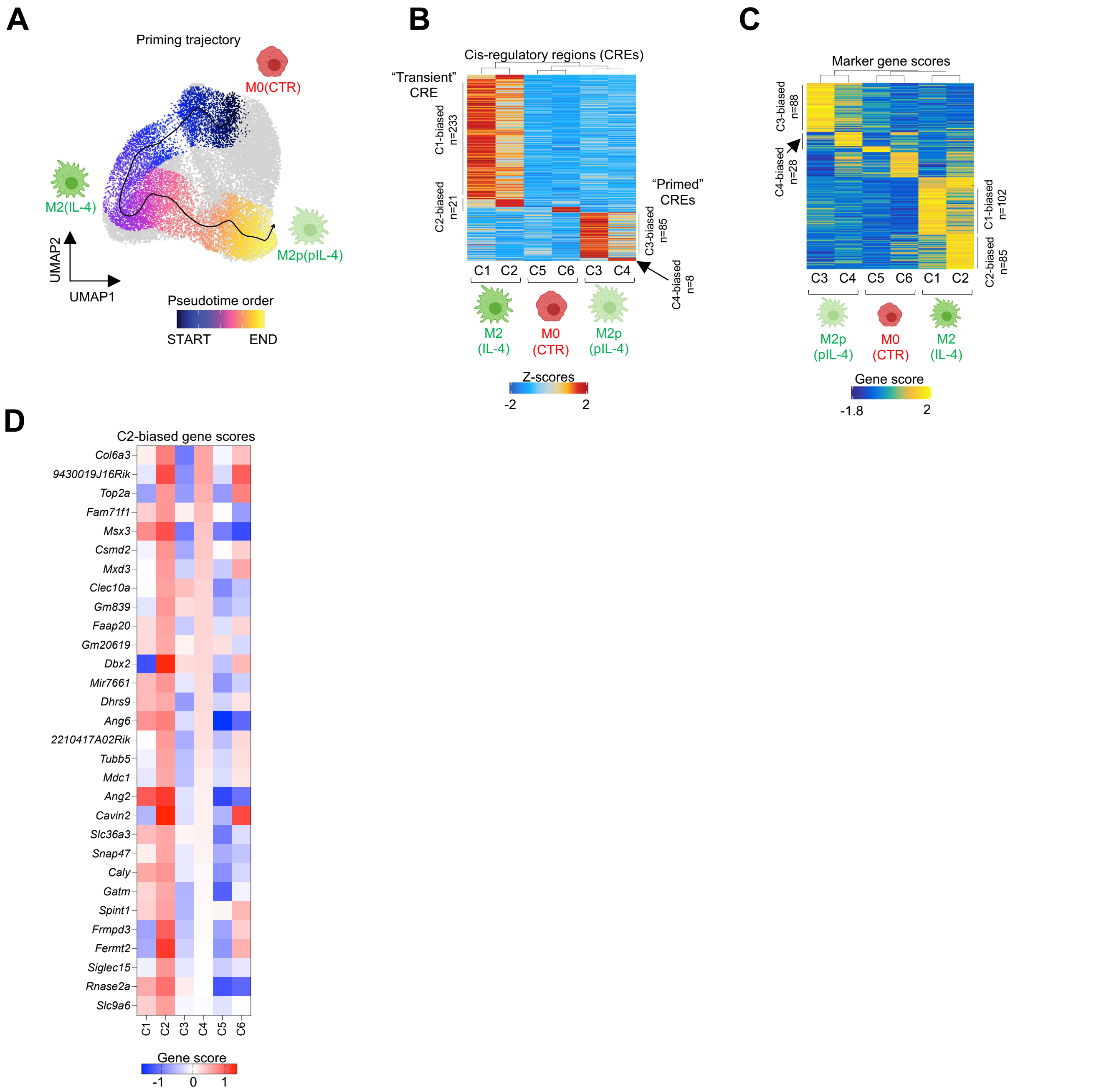

### Supplementary Figure 6.

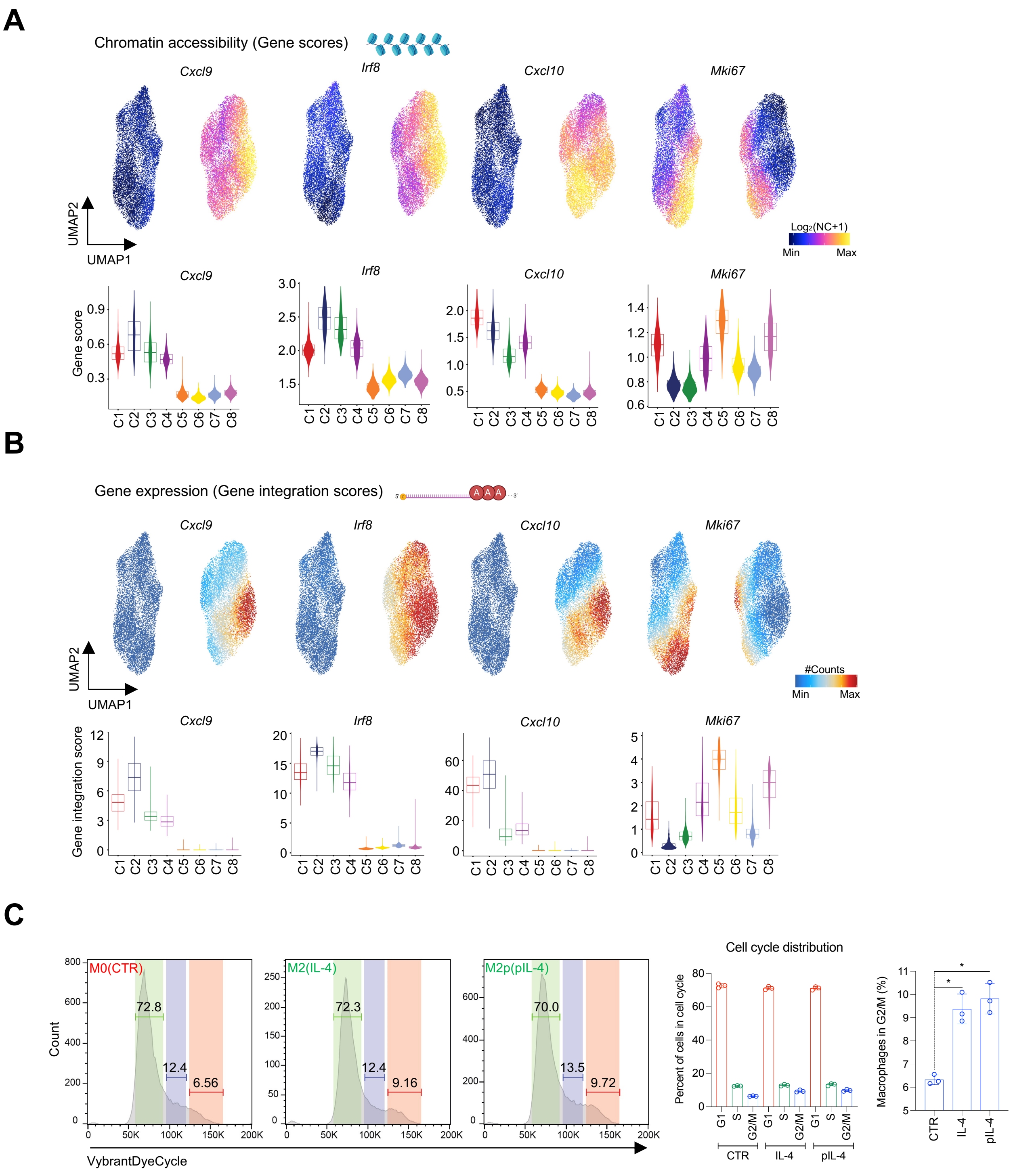

### Supplementary Figure 7.

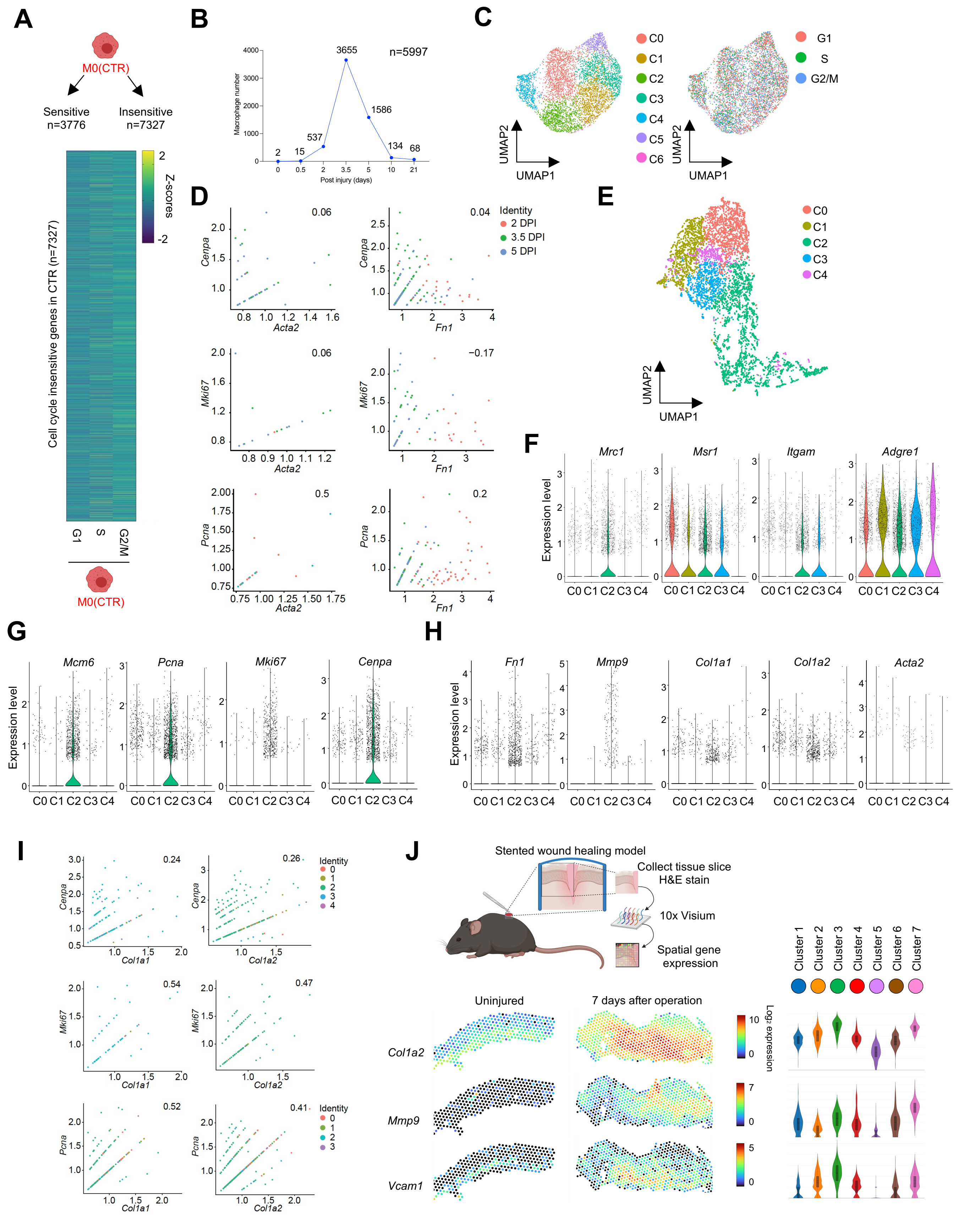
